# The small non-coding RNA B11 regulates multiple facets of *Mycobacterium abscessus* virulence

**DOI:** 10.1101/2022.10.27.514065

**Authors:** Michal Bar-Oz, Maria Carla Martini, Maria Natalia Alonso, Michal Meir, Nicola Ivan Lore, Paolo Miotto, Camilla Riva, Junpei Xiao, Catherine S. Masiello, Maria-Anna Misiakou, Huaming Sun, Justin K. Moy, Helle Krogh Johansen, Daniela Maria Cirillo, Scarlet S. Shell, Daniel Barkan

**Author notes:** Equal contributions (Michal Bar-Oz, Maria Carla Martini, Maria Natalia Alonso).

## Abstract

*Mycobacterium abscessus* causes severe, virtually incurable disease in young patients with cystic fibrosis. Little is known in *M. abscessus* about the roles of small regulatory RNAs (sRNA) in gene expression regulation. Here, we show that the sRNA B11 controls gene expression and virulence-associated phenotypes in this pathogen. B11 deletion from the smooth strain ATCC_19977 produced a rough colony morphology, increased pro-inflammatory signaling and virulence in *in-vivo* infection models, and increased resistance to clinically relevant antibiotics. Examination of clinical isolate cohorts revealed some isolates with B11 mutations or reduced expression. We used RNAseq and proteomics to investigate the effects of B11 on gene expression and test the impact of two mutations found in clinical isolates. Approximate 230 genes were differentially expressed in the B11 deletion mutant. Strains with the clinical B11 mutations showed similar expression trends to the deletion mutant but of a lesser magnitude, suggesting partial loss of function. Among genes upregulated in the B11 mutant, there was a strong enrichment for genes with B11-complementary sequences in their predicted ribosome binding sites (RBS), consistent with a model of translational repression via base-pairing of B11 to RBSs. Comparing the proteomes similarly revealed that upregulated proteins were strongly enriched for B11-complementary sequences in their RBS, consistent with B11 functioning as a negative regulator through direct binding of target mRNAs. Intriguingly, the genes upregulated in the absence of B11 included components of the ESX-4 secretion system, known to be critical for *M. abscessus* virulence. One of these genes had a B11-complementary sequence at its RBS, and fusing the UTR of this gene to a reporter was sufficient to make the reporter suppressible by B11. Taken together, our data show that B11 may act as either a negative or positive regulator with pleiotropic effects on gene expression and clinically important phenotypes in *M. abscessus*. The presence of hypomorphic B11 mutations in clinical strains supports the idea that lower B11 activity may be advantageous for *M. abscessus* in some clinical contexts. To our knowledge, this is the first report of the role of an sRNA in *M. abscessus*.

## INTRODUCTION

Non-tuberculous mycobacteria are increasingly recognized as human pathogens (1–4). Of these, *Mycobacterium abscessus* is associated with the most severe chronic infections, including virtually incurable pulmonary disease in patients with cystic fibrosis (CF) (5–7). Bacterial virulence requires regulation of gene expression to facilitate phenotypic adaptation to various niches within the human host. However, knowledge of the fundamental biology underlying gene regulation and phenotypic adaptations in *M. abscessus* lags far behind that of *M. tuberculosis* (Mtb), in part due to lack of genetic tools and in part due to the relatively recent recognition of *M. abscessus* as an important human pathogen.

Colony morphology is often considered a correlate of *M. abscessus* virulence. Patients are typically infected with smooth (S) variants. The S morphotype is conferred by the presence of glycopeptidolipids (GPL) in the cell envelope. Rough (R) variants, not producing GPL, typically arise later in infection (8,9). S and R morphotypes exhibit different growth characteristics in macrophages (10), and different stimulation of innate immunity (11–14). Overall, S morphotypes are typically associated with intracellular growth and blockage of phagolysosomal fusion while R morphotypes are associated with extracellular growth and pro-inflammatory signaling that causes severe tissue destruction (9,10,15–17). However, opposite effects have been shown with respect to intracellular growth highlighting the complexity of this phenotype (11). The switch is usually due to mutations in *gpl* biosynthesis genes such as mps1 and mps2 (*MAB_4099c*, *MAB_4098c*, respectively) or GPL transporter genes such as *mmpL4b (MAB_4115c)* (18,19).

As in Mtb, ESX secretion systems are present in *M. abscessus* and involved in virulence. The ESX-4 system was shown to be important for survival of *M. abscessus* in macrophages and may play a role in phagosomal permeabilization, functionally analogous to the role of ESX-1 in Mtb (20). However, little is known about how expression and function of the ESX-4 system and other *M. abscessus* virulence factors are regulated.

Small non-coding RNAs (sRNA) contribute to pathogenesis and regulation of virulence in bacteria including *Listeria monocytogenes* (21), *Staphylococcus aureus* (22), and *Salmonella enterica* (23,24). sRNAs have been identified in mycobacteria, mostly by RNAseq screens (25–28). Some are expressed in specific conditions, giving clues regarding their roles (29). The *M. smegmatis* sRNA Ms1 is upregulated in stationary phase and interacts with core RNA polymerase, repressing transcription by preventing holoenzyme formation (30). Consistently, the Mtb homolog MTS2823 is highly expressed in stationary phase, and ectopic overexpression in log phase caused downregulation of energy metabolism genes (26). Mcr11 is involved in regulation of fatty acid metabolism in Mtb (31), whereas Mcr7 is involved in TAT secretion (32). MrsI is induced by multiple infection-associated stressors and required for mycobacterial growth in low-iron conditions (33). MTS1338 was suggested to trigger dormancy-like gene expression changes in response to infection-associated conditions in Mtb (34) as well as to mediate response to oxidative stress (35). The Mtb sRNA B11 was reported to cause slow growth and repress several genes when overexpressed in *M. smegmatis*, due in some cases to base-pairing of B11 to genes’ ribosome binding sites (RBSs) and presumably competing with ribosomes (25,36). Orthologs of some of these sRNAs were identified in a transcriptomics study of *M. abscessus* (37). However, almost nothing is known about roles played by sRNAs in this important emerging pathogen.

Here we screened an *M. abscessus* transposon mutant library and identified the sRNA B11 as a factor required for smooth morphology. We constructed a targeted B11 deletion mutant and characterized it for gene expression and pathogenesis-associated phenotypes. We found that loss of B11 caused increased virulence and pro-inflammatory immune signaling as well as altered expression of a large set of genes enriched for those with B11-complementary sequences in their RBSs. Genes with increased expression in the B11 deletion mutant included several components of the ESX-4 secretion system. Furthermore, B11 is mutated in several clinical isolates. Strains with these mutations have gene expression patterns consistent with reduced B11 activity, suggesting that reduced B11 function is advantageous to the bacteria in some clinical contexts.

## RESULTS

### Loss of the sRNA B11 causes rough colony morphology in *M. abscessus*

While mutations in GPL biosynthesis and transport genes are known to cause conversion of smooth *M. abscessus* to rough, we sought to determine if there were additional genetic paths to the rough morphotype. We therefore constructed a transposon mutant library in *M. abscessus* ATCC_19977 (S morphotype) (38) and screened it for rough colonies. Some rough colonies had transposons inserted in GPL biosynthesis and transport genes (*mps1*, *mps2*, and *mmpL4b*), as expected (18). We also identified a rough mutant with a transposon inserted in the −10 sigma factor binding site of the promoter for the sRNA B11 (Fig. 1a and S1a-b). The Tn mutant had substantially less B11 transcript than the parental strain (Fig. 1b, WT and Tn-B11). Mycobacterial promoters typically contain the core −10 sequence TANNNT (39–41). As the Tn left this core sequence intact, B11 was likely expressed from a fusion of its native promoter and the 3’ portion of the Tn.

**Figure 1.**
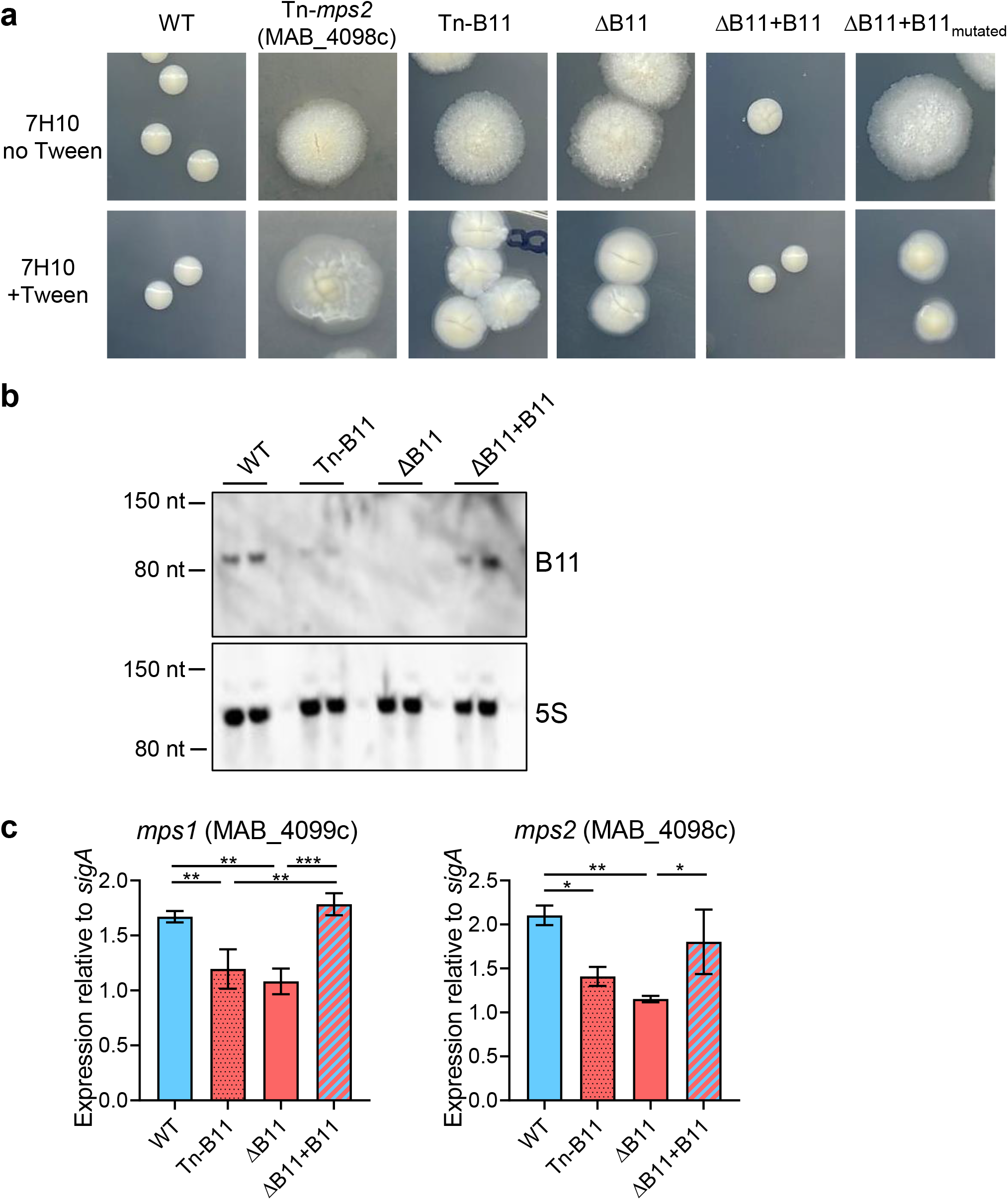
Disruption of the sRNA B11 causes rough morphology in *M. abscessus*. **A** The indicated strains were grown in liquid media with Tween 80, diluted and plated on soli media with and without Tween 80. Colonies were imaged after 7 days of growth. **B**. Norther blotting of duplicate RNA samples from the indicated strains confirmed that B11 expressio was reduced in the transposon mutant, absent from the deletion mutant, and restored in th complemented mutant. **C**. Quantitative PCR reveals that expression of two key GP biosynthesis genes is modestly but significantly reduced in strains lacking B11. * *P* < 0.05; * *P* < 0.01; *** *P* < 0.001; ANOVA with Tukey post-test. qPCR comparisons where *P* > 0.0 are not shown. qPCR data represent triplicate log-phase cultures.

To determine if reduced B11 expression was responsible for the rough morphology of the Tn mutant, we replaced the native B11 gene in ATCC_19977 with a *zeocin^R^* gene. The ΔB11 mutant had characteristic rough morphology and exhibited extensive clumping in liquid culture, especially in media without Tween (Fig. 1a and S1c-d). Both phenotypes reverted to WT in a complemented mutant with one copy of B11 transcribed from the native promoter, integrated at the L5 site (ΔB11+B11) (Fig. 1a-b). However, the colony morphology was not complemented by a version of B11 with multiple mutations in one of the unstructured loops (Fig. 1a and Fig. S1e), implicating the loop as playing a functionally important role.

To investigate the mechanistic basis of the rough morphology of ΔB11, we measured expression of two key GPL biosynthesis genes, *mps1* (*MAB_0499c*) and *mps2* (*MAB_0498c*). Both had significantly reduced expression in the ΔB11 mutant and returned to WT levels upon complementation (Fig. 1c), suggesting the rough morphology was due in part to reduced expression of GPL biosynthesis genes.

### A B11 deletion strain has virulence and envelope-leakiness properties reminiscent of “classic”, GPL-related rough strains

Rough *M. abscessus* strains have been reported to be more virulent and more pro-inflammatory. We therefore wondered if the ΔB11 mutant differed from its smooth WT parental strain with respect to these properties. To assess virulence, we first used our previously established *Galleria mellonella* infection model (42). We infected 60 larvae with WT *M. abscessus* (ATCC_19977) or the ΔB11 mutant, using an inoculum of the ΔB11 mutant approximately half that of WT to ensure that we could have confidence in any observed hypervirulence phenotype in the mutant. Despite the lower initial inoculum, larval survival was significantly reduced in the ΔB11-infected group (Fig. 2a). Hypervirulence is often associated with higher bacterial loads. However, bacterial loads were not higher in the larva infected by the ΔB11 mutant when measured at an early timepoint (Fig. 2b), despite significantly lower survival.

**Figure 2.**
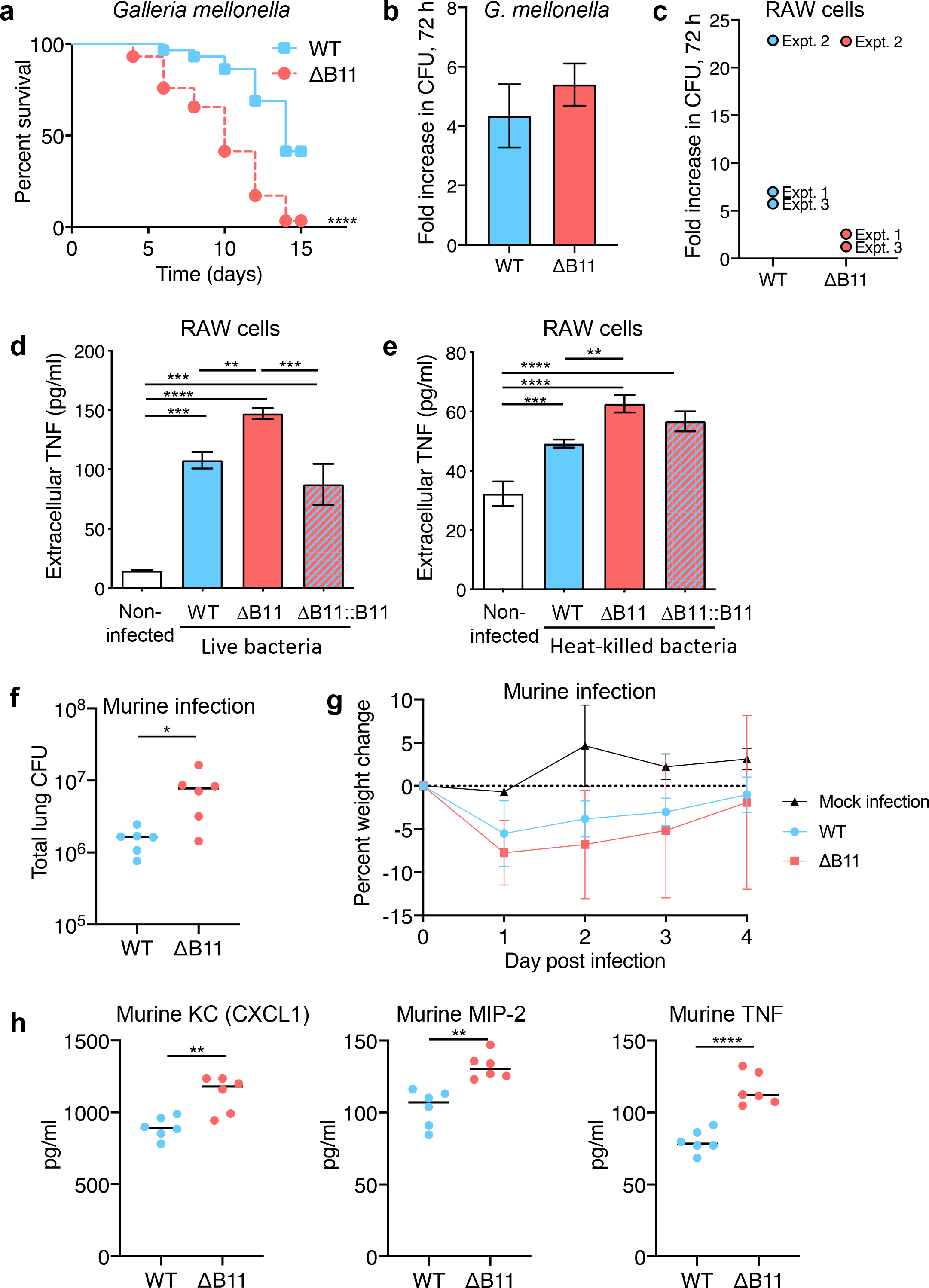
Disruption of *M. abscessus* B11 increases virulence and cytokine secretion. **A**. *Galleria mellonella* infections. 30 larvae per group were infected by either ~1000 CFU of WT *M. abscessus* or ~500 CFU of ΔB11 mutant. Larvae were followed for two weeks. The survival in each group is showed in a Kaplan-Mayer curve. ****, *P* < 0.0001 for the comparison between strains. **B**. Bacteria from infected *G. mellonella* were enumerated three days after infection and no significant difference between strains was found. **C**. RAW macrophage-like cells were infected with the indicated *M. abscessus* strains and bacteria were enumerated three days later. Three separate experiments were performed (expt. 1-expt. 3) with 2-4 replicates per experiment. The mean fold-increase for each experiment is shown. **D**. RAW cells were infected with *M. abscessus* at an MOI of 5 and TNF secretion was measured after 6 hours. **E**. RAW cells were exposed to heat-killed *M. abscessus* at the equivalent to an MOI of 20, washed, and TNF secretion measured after 24 hours. **F**. Lung bacterial burden for C57BL/6NCrl mice four days after infection with the indicated *M. abscessus* strains. **G**. Weight change in mice following infection with the indicated *M. abscessus* strains or mock infection with PBS-infused agar beads. Points are error bars represent mean and SD. n=9 mice per group for infections and 3 mice for the mock infrection. **H**. The indicated cytokines were measured in lung homogenates four days after infection with the indicated *M. abscessus* strains. Bar charts display means and SD. ** *P* < 0.01; *** *P* < 0.001; **** *P* < 0.0001, ANOVA with Tukey post-test.

To assess the effects of B11 on intracellular replication and innate immune activation, we infected RAW mouse macrophage-like cells with ATCC_19977 or the ΔB11 mutant. Proliferation of the ΔB11 mutant after 72 hours was comparable to that of WT (Fig. 2c). However, RAW cells infected with the ΔB11 mutant secreted more TNF than those infected by the WT or complemented strains (Fig. 2d), suggesting stronger activation of an inflammatory response (43). RAW cells exposed to heat-killed ΔB11 bacteria also secreted more TNF than those exposed to heat-killed WT bacteria (Fig. 2e), indicating that this response may result directly from the modified cell envelope of the deletion mutant.

To determine if the hyper-inflammatory phenotype was preserved in a whole animal experimental model, we infected mice with the WT and ΔB11 strains, using our established agar-beads infection model (44,45). Bacterial burden after four days of infection was significantly higher for mice infected with ΔB11 compared to those infected with wildtype (Fig. 2f). Similarly, mice infected with ΔB11 exhibited a trend towards more weight loss (Fig. 2g) and had substantially higher lung levels of the pro-inflammatory cytokines TNF, Keratinocyte-derived Cytokine (KC, also known as CXCL1), and Macrophage-Inflammatory Protein-2 (MIP-2, also known as CXCL2) (Fig. 2h).

To further compare the cell-envelope-related properties of the ΔB11 strain, we examined the proteins released into culture supernatants. The ΔB11 strain released substantially more proteins of various sizes compared to the WT and complemented strains (Fig. S2a). A “classic” rough strain in which the GPL biosynthesis gene *mps2* was disrupted by a transposon (Tn-*mps2*) similarly released more proteins into the culture supernatant, suggesting this may be a general property of rough strains (Fig. S2b). To determine if the greater protein abundance in the culture supernatants was due exclusively to increased secretion, we transformed the strains with a plasmid encoding cytoplasmic mCherry and probed the culture supernatants and cell lysates by western blot. In the WT strain mCherry was detected only in cell lysates, while in the ΔB11 and Tn-*mps2* strains it was detected in the culture supernatants as well (Fig. S2c). As mCherry does not contain a secretion signal sequence, this result suggests that rough strains may be generally leakier, with a greater amount of cytoplasmic protein being released during normal growth.

### B11 affects drug resistance

The ΔB11 mutant was also more resistant to two antimycobacterial drugs. The MICs of linezolid and rifampicin were elevated 4-fold and 8-fold, respectively. In contrast, only a trivial difference was found in the MIC of ciprofloxacin (Table 1), and no change was found in the MICs of meropenem, amikacin, azithromycin, or tigecycline. To our knowledge rough morphology *per se* has not been reported to affect drug resistance, specifically not at these concentrations; these effects may therefore be due to other physiological changes in the ΔB11 mutant.

**Table 1.** *M. abscessus* ΔB11 is more resistant to linezolid and rifampin as compared to its WT and complemented (ΔB11+B11) counterparts.

|  |  | **WT *M. abscessus*<sup>Lux</sup> (ATCC 19977)** | ***M. abscessus*<sup>Lux</sup> ΔB11** | ***M. abscessus*<sup>Lux</sup> ΔB11+B11** |
| --- | --- | --- | --- | --- |
| **Ciprofloxacin** | MIC<sub>90</sub> | 0.3 µg/ml | 0.3 µg/ml | 0.3 µg/ml |
|  | MIC<sub>99</sub> | 0.6 µg/ml | 0.6 µg/ml | 0.6 µg/ml |
| **Linezolid** | MIC<sub>90</sub> | 0.8 µg/ml | 1.6 µg/ml | 0.8 µg/ml |
|  | MIC<sub>99</sub> | 1.6 µg/ml | 12.5 µg/ml | 1.6 µg/ml |
| **Rifampin** | MIC<sub>90</sub> | 2.35 µg/ml | 4.7 µg/ml | 4.7 µg/ml |
|  | MIC<sub>99</sub> | 4.7 µg/ml | 37.5 µg/ml | 9.4 µg/ml |

### B11 affects expression of a large gene set enriched for genes with B11-complementary sequences in their 5’ UTRs

Heterologous overexpression of *M. tuberculosis* B11 in *M. smegmatis* was previously shown to affect gene expression, in some cases through direct binding to RBSs (36). To determine the effect of B11 on gene expression in *M. abscessus*, we performed whole-proteome analysis and RNAseq (Table S1). RNAseq with the WT, ΔB11, and ΔB11+B11 strains revealed 252 genes that were differentially expressed in the deletion strain compared to the WT strain (Fig 3a, fold change >= 2 and adjusted *p* < 0.05) and 229 that were differentially expressed in the deletion strain compared to the complemented strain (Fig 3b). LC-MS/MS of whole cell lysates from WT and ΔB11 strains revealed 90 differentially abundant proteins (Fig 3c, proteins with >= 2-fold expression change relative to WT and CVs <0.5 between replicates for each strain). More genes were upregulated than downregulated in the ΔB11 strain in both the proteomics and RNAseq datasets, consistent with previous postulation that B11 is a negative regulator in some mycobacteria (36). There was a significant correlation between expression changes in the ΔB11 strain vs. WT in the proteomics and RNAseq datasets (Fig 3d).

**Figure 3.**
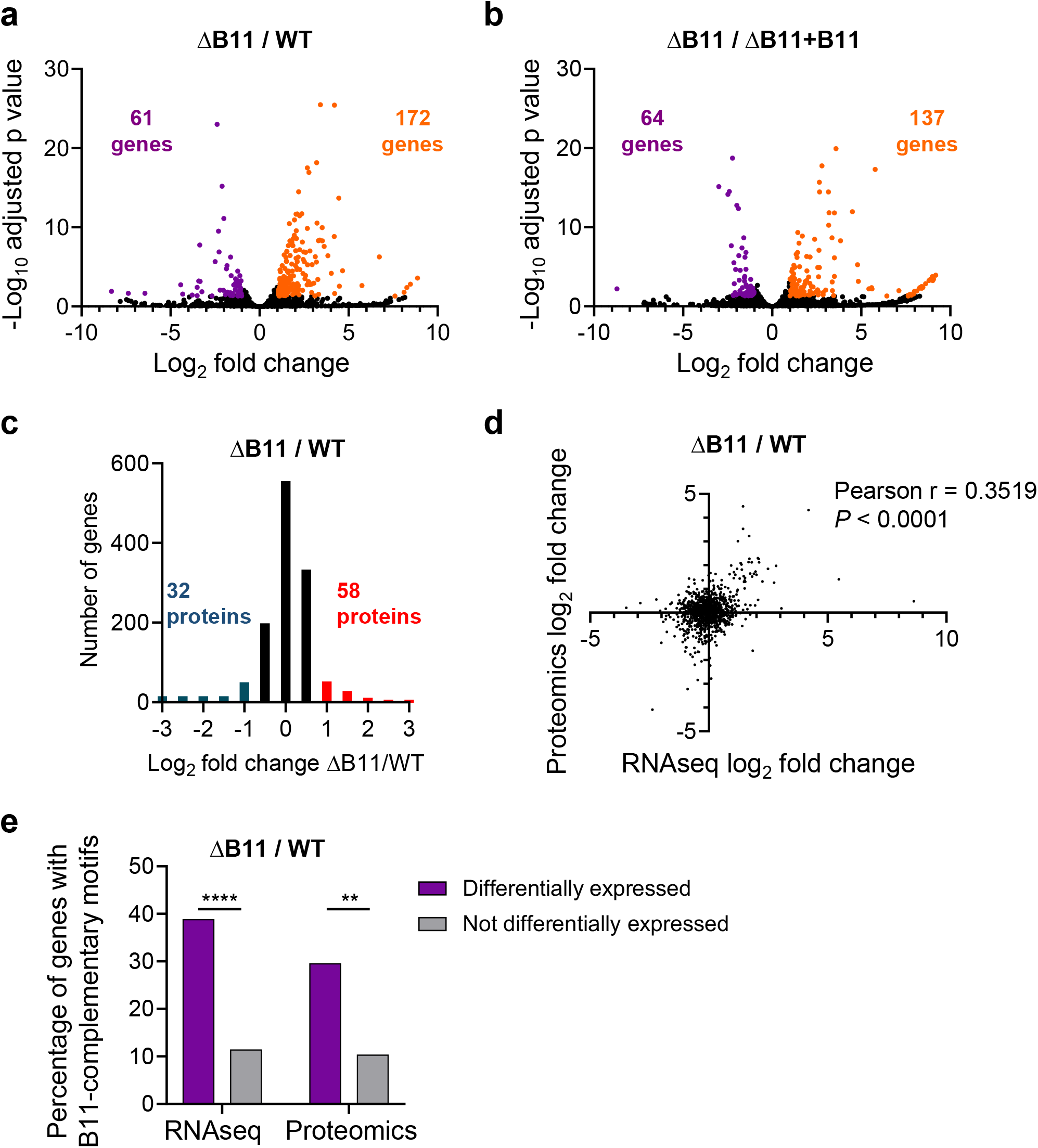
Disruption of the sRNA B11 in *M. abscessus* causes gene expression chan at the mRNA and protein levels particularly in genes with B11-complementary riboso binding sites. **A-B**. Volcano plots showing gene expression changes in log-phase when was deleted. Genes that met the thresholds of >= 2-fold expression change and adjusted 0.05 are indicated by orange and purple. **C**. LC-MS/MS to compare protein abundance in WT and ΔB11 strains. Proteins with >= 2-fold abundance differences and between-replicate < 0.5 are indicated by red and blue. **D**. Protein and mRNA-level expression changes w correlated. **E**. Differentially expressed genes are enriched for the presence of B complementary sequences in their ribosome bind sites. Only genes (and their encoded prote with defined transcription start sites were included in the analysis. Genes and proteins w classified as differentially expressed if they met the criteria indicated above. ** *P* < 0.01; **** 0.0001, Fisher’s exact test.

To investigate the possibility that some of the expression changes were due to direct base-pairing of B11 to RBSs, we quantified the presence of B11-complementary sequences in the 25 nt upstream of the start codons of genes with defined 5’ UTRs (37) (Table S1). In the proteomics and RNAseq datasets respectively, 30 and 37% of differentially expressed genes with defined 5’ UTRs had B11-complementary sequences of 6 nt or greater, compared to 10-11% of non-differentially-expressed genes (Fig 3e). This represented a substantial enrichment for B11-complementary RBS sequences among the differentially expressed genes. This is consistent with the ideas that (i) B11 may directly repress translation (36) and (ii) reduced translation or ribosome binding frequently lead to mRNA degradation and/or premature transcriptional termination in bacteria (46–52) and reviewed in (53).

Genes affected by B11 deletion that do not have B11-complementary regions in their 5’ UTRs may be affected indirectly by transcription factors that are in turn directly or indirectly affected by B11.

### B11 represses expression of several ESX-related genes

The genes differentially expressed by RNAseq included all eight genes reported to encode components of the ESX-4 secretion system and some possible ESX regulators (20). Most of these either had B11-complementary sequences in their RBSs, or were in operons downstream of genes that had B11-complemantary sequence in their RBSs. In one case, the gene was leaderless with a B11-complementary sequence early in the coding sequence (Fig. S3). To validate some of these expression changes, we performed qPCR on RNA from independently grown cultures. In most cases, deletion of B11 led to increased expression that was complemented when B11 was ectopically expressed (Fig. 4a-b). A version of B11 with extensive mutations in one of the loops (Fig. S1e) did not complement expression in most cases where it was tested (Fig. 4b).

**Figure 4.**
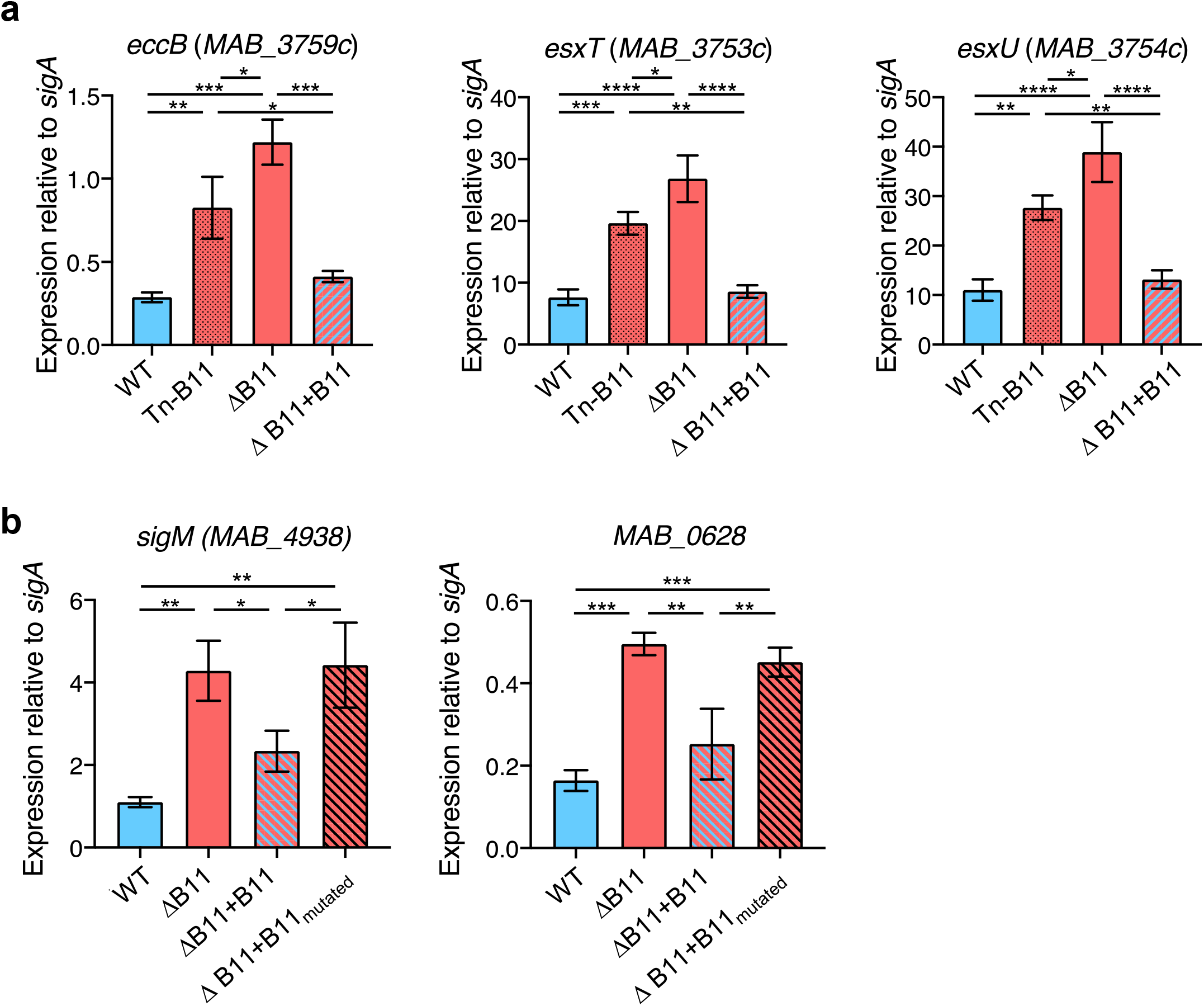
Disruption of *M. abscessus* B11 causes increased expression of several ESX-related genes with B11-complementary sequences. **A**. Disruption or deletion of B11 causes increased expression of genes encoding the EsxU, EsxT, and the ESX-4 structural protein EccB. **B**. Deletion of B11 causes increased expression of *sigM* and *MAB_0628*, which is predicted to be transcribed upstream of *espI* on a polycistronic transcript (see Fig. S3). ΔB11+B11_mutated_ denotes complementation with a B11 variant harboring the mutations indicated by blue squares in Fig. S1e. Bars display means and SD. * *P* < 0.05; ** *P* < 0.01; *** *P* < 0.001; **** *P* < 0.0001, ANOVA with Tukey post-test. qPCR comparisons where *P* > 0.05 are not shown. qPCR data represent triplicate log-phase cultures.

### B11 represses expression of a critical component of the ESX-4 secretion system via its unstructured loops

One of the most strongly upregulated genes in the ΔB11 strain was *eccB4* (*MAB_3759c*), which was upregulated at both the mRNA and protein levels. EccB4 is one of the structural components of the ESX-4 secretion system and is required for *M. abscessus* growth in macrophages (20). The *eccB4* 5’ UTR has a 7-nt sequence that is complementary to loop 2 of B11, as well as to 6 nt of loop 1 (Fig 5a). To test the hypothesis that B11 negatively regulates *eccB4* by binding to its 5’ UTR, we constructed a set of reporters in which mCherry was expressed from the synthetic MOP promoter either with the MOP-associated synthetic 5’ UTR or with the *eccB4* 5’ UTR (Fig 5b-c). mCherry expression from the construct with the MOP 5’ UTR was similar for the WT and ΔB11 strains (Fig 5b). In contrast, mCherry expression from the construct with the *eccB4* 5’ UTR was substantially higher in the ΔB11 strain compared to the WT and complemented strains. The *eccB4* 5’ UTR contains a tract of six guanosines that could theoretically bind to either of B11’s two C-rich loops. To determine which loop was required for repression of expression via the *eccB4* 5’ UTR, we mutated each of the two loops individually and in combination (Fig 5b and d). B11 was still able to repress mCherry expression when either loop was mutated individually, but variants with mutations in both loops could not, strongly suggesting these loops play a direct role in gene regulation by B11.

**Figure 5.**
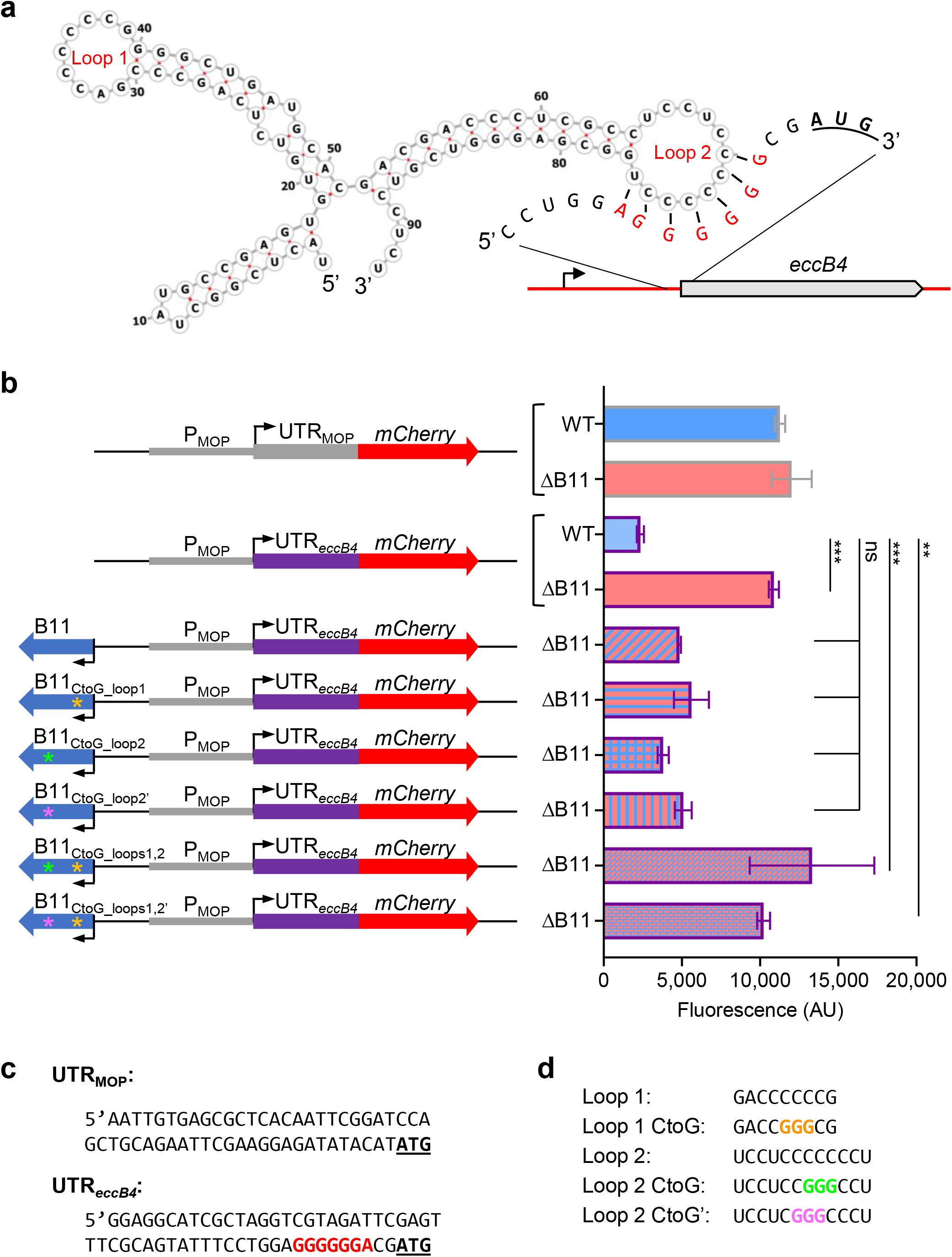
The 5’ UTR of *eccB4* is sufficient to make a reporter gene B11-repressible in *M. abscessus*, and either of B11’s unstructured loops are sufficient for such repression. **A**. Schematic of the predicted secondary structure of B11 demonstrating the possibility of base-pairing to the ribosome binding site of the *eccB4* 5’ UTR. The *eccB4* start codon is underlined and bolded. Note that the 6-G sequence in the UTR could also base-pair with loop 1 of B11. **B**. The constructs shown schematically on the left were transformed into WT or ΔB11 *M. abscessus* as indicated and fluorescence was measured by flow cytometry of biological replicate cultures. The WT strain harboring the *eccB4* UTR-mCherry construct was compared to the strains below by ANOVA followed by Dunnett’s multiple comparisons test. Bar chart displays means and SD. ** *P* < 0.01; *** *P* < 0.001. **C**. The complete sequences of the two 5’ UTRs used in the constructs in B. **D**. The sequences of the B11 loops and the mutations used in B.

To test if the tract of six guanosines present in *eccB4*’s UTR is needed for the direct interaction with B11’s cytosine-loops, we constructed another reporter system. In this system, the red fluorophore mCherry (under a B11-unresponsive promoter and UTR, as above) and the green fluorophore mWasabi (under the same B11-neutral promoter, but with the UTR taken from *eccB4*, including the six guanosines) were placed on the same, single copy integrating plasmid, and the relative fluorescence of mWasabi was compared to the constant fluorescence of mCherry in each bacterium by flow cytometry. As seen in Fig. 6, the ratio of green:red fluorescence increased substantially in the ΔB11 mutant as compared to the ratio in WT, indicating the deletion of B11 de-repressed the *eccB4* UTR. However, when a similar experiment was performed where the tract of six guanosines was interrupted by mutating it to 5’-GTGTGT-3’, the increase in the ratio was no longer present, as the UTR, now unable to bind B11, was already de-repressed in the WT. Taken together, these data indicate (i) that the 5’ UTR of *eccB4* is sufficient to make a transcript B11-suppressible, (ii) that the guanosine tract in the UTR is necessary for such repression, and (iii) that either of B11’s two C-rich loops are sufficient to carry out this repression.

**Figure 6.**
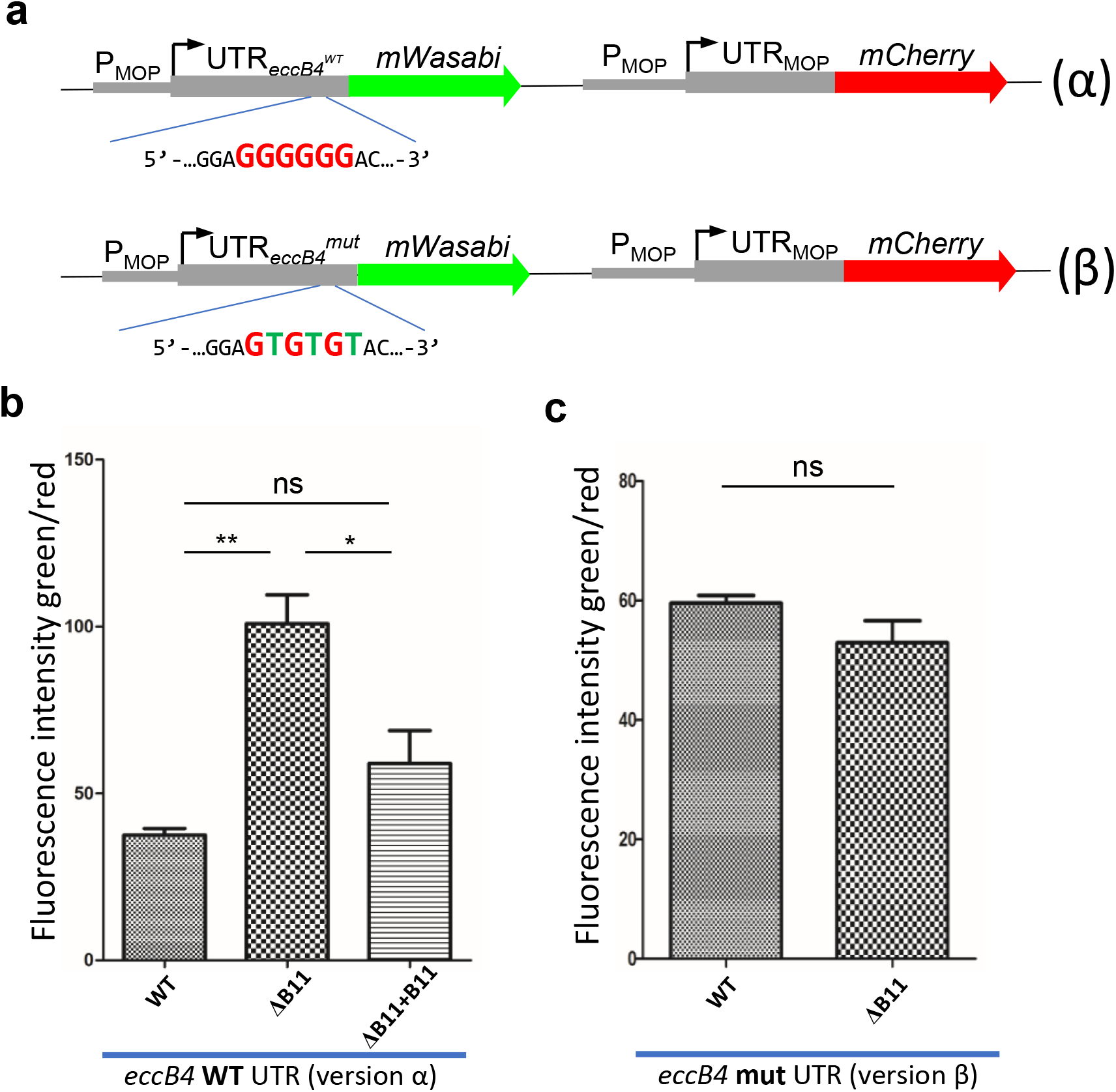
The string of six guanidines in *eccB4*’s 5’ UTR is essential for repression by the sRNA B11. **A**. two versions of an otherwise similar plasmid were constructed, where the fluorophore mCherry is under a B11-neutral promoter and UTR (with constitutive expression), whereas mWasabi is under the same promoter, with the UTR of *eccB4*, containing a string of guanosines (α version), or where the string of guanosines was mutated (β version). **B**. The intensity of the green (wasabi-originating) fluorescence relative to the red (mCherry) fluorescence was measured by flow cytometry in WT, ΔB11, and complemented bacteria carrying the WT (α) version of the plasmid. * *P* < 0.05; ** *P*< 0.01, ANOVA with Tukey post-test. **C**. The intensity of the green relative to red fluorescence was measured by flow cytometry in WT and ΔB11 bacteria carrying the mutated (β) version of the plasmid. ns *P* > 0.05, unpaired t test. Error bars represent SEM for both plots.

### Loss or reduction of B11 function occurs in some clinical M. abscessus isolates

Given the association between rough morphotypes and *M. abscessus* disease progression, we wondered if downregulation of B11 expression or activity occurred in clinical isolates as part of an *in-vivo* evolutionary process favoring progression to more virulent phenotypes. We examined a recently published cohort of 70 Irish *M. abscessus* clinical isolates that underwent whole genome sequencing (54). Two of the seventy isolates had indel mutations in the B11 loops: one deletion of a C in the first loop (Shortening it from 6 Cs to 5 Cs; B11_del1_), and one insertion of a C in the second loop (from 7 Cs to 8; B11_ins1_; Fig 7a). We also examined a set of 52 clinical isolates from Denmark and found that two had the same B11_ins1_ mutation. To test the hypothesis that these mutations affect B11 stability or function, we complemented the ΔB11 strain with B11 harboring the deletion in loop 1 or the insertion in loop 2. Both variants had similar abundance to WT B11 (Fig S4), suggesting the mutations do not affect B11 expression or stability.

**Figure 7.**
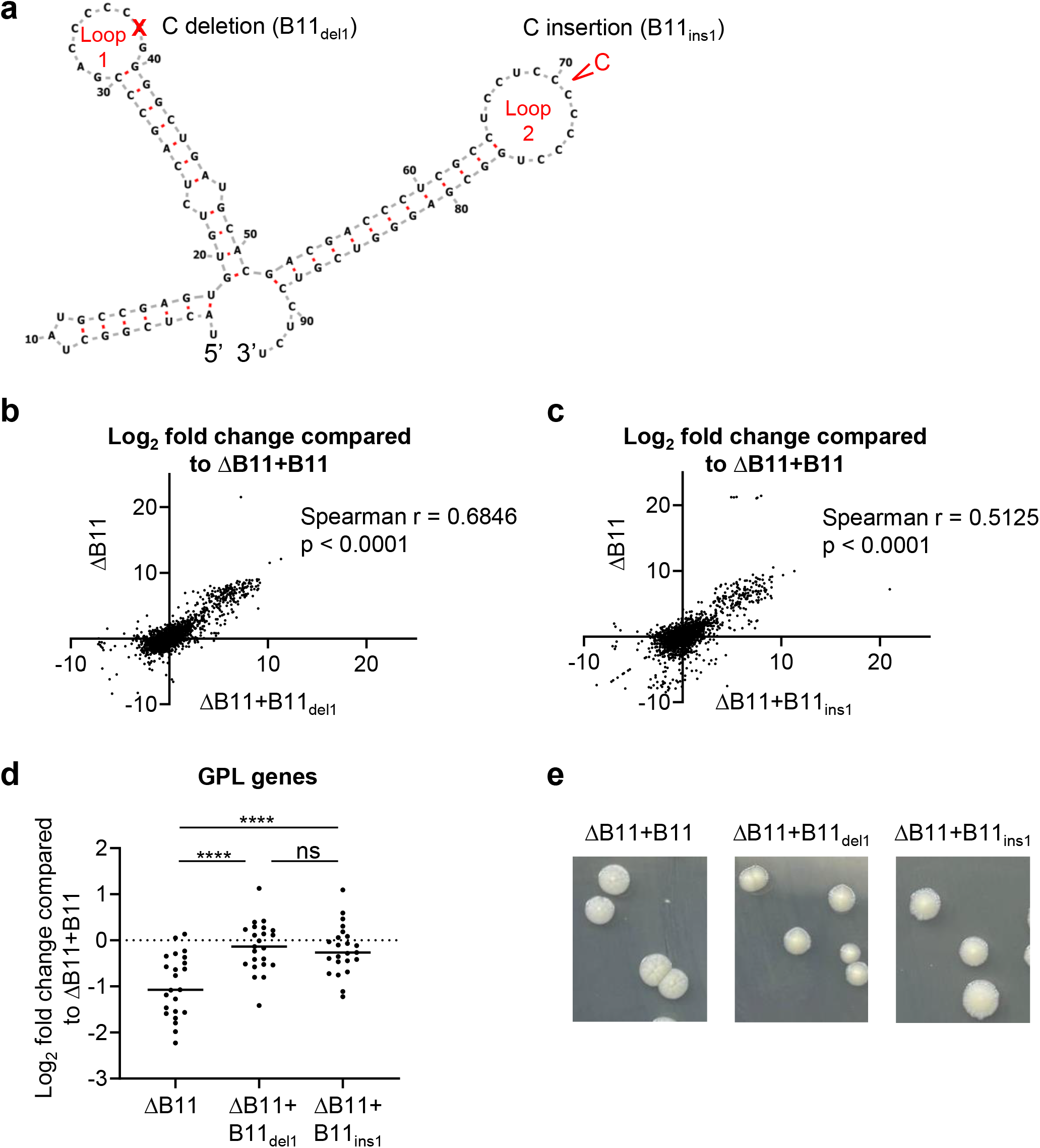
B11 mutations found in clinical *M. abscessus* strains cause partial loss of function. **A**. Deletion of one C from loop 1 was found in one clinical isolate, while insertion of one C into loop 2 was found in 3 clinical isolates. **B-C**. Transcriptome-wide RNAseq profiles of the ΔB11 strain alone or harboring each of the two clinical loop mutations were correlated when compared to the ΔB11 strain transformed with WT B11. **D**. RNAseq data revealed that genes involved in GPL synthesis and transport had reduced expression in the ΔB11 strain and this was complemented by each of the clinical loop mutations. **** *P* < 0.0001, one way RM ANOVA. **E**. Smooth morphology was largely restored upon complementation of the ΔB11 strain with B11 harboring each of the two clinical loop mutations

To assess the functional impact of the two clinical loop mutations, we performed RNAseq on the ΔB11 strain complemented with each mutated version of B11 (Table S1). Strikingly, the gene expression changes in each loop mutant were strongly correlated with the expression changes in the ΔB11 strain (Fig 7b-c; Spearman r = 0.68 and 0.51 for ΔB11 vs. ΔB11+B11_del1_ and ΔB11 vs ΔB11+B11_ins1_ respectively, *p* < 0.0001 for both). In this analysis we compared the ΔB11 strain and loop mutant strains to the ΔB11+B11 strain rather than the WT strain, to ensure that the results were not confounded by changes due to expression of B11 from an ectopic location. These correlations revealed that many of the expression changes caused by loss of B11 were also present in the loop mutants, albeit to a lesser extent, thus in some cases not meeting our strict criteria for differential expression (fold-change >= 2 and *p* < 0.05; 83 genes for ΔB11+B11_ins1_/ΔB11+B11 and 20 genes ΔB11+B11_del1_/ΔB11+B11). The two clinical mutations therefore appear to be hypomorphic, causing partial loss of function.

Interestingly, GPL biosynthesis and transport genes, which were underexpressed in the ΔB11 strain, were largely restored by both B11 loop mutants (Fig. 7d, Table S2, (55)), consistent with the observation that both variants largely restored smooth morphology, with subtle differences compared to WT B11 (Fig 7e).

Finally, we examined B11 expression in a set of clinical isolates from Israel, and found that one isolate, termed LAH, had a dramatic reduction in B11 abundance compared to ATCC_19977 and the other clinical strains (Fig 8a). Sequencing of B11 and the surrounding region did not reveal any mutations that could explain why B11 has reduced expression in LAH, suggesting *trans*-regulatory changes. To assess the functional impact of the absence of B11 in this strain, we transformed it with a plasmid to ectopically express B11 from a constitutive artificial promoter (MOP) and measured resistance to linezolid and rifampicin. Consistent with our finding that deletion of B11 from ATCC_19977 increased resistance to these antibiotics, ectopic expression of B11 caused increased sensitivity in LAH (Fig 8b). Taken together, our findings suggest that mutations and regulatory changes that either abrogate the expression, or affect the biologic potency of the B11 transcript, are found in clinical isolates throughout the globe.

**Figure 8.**
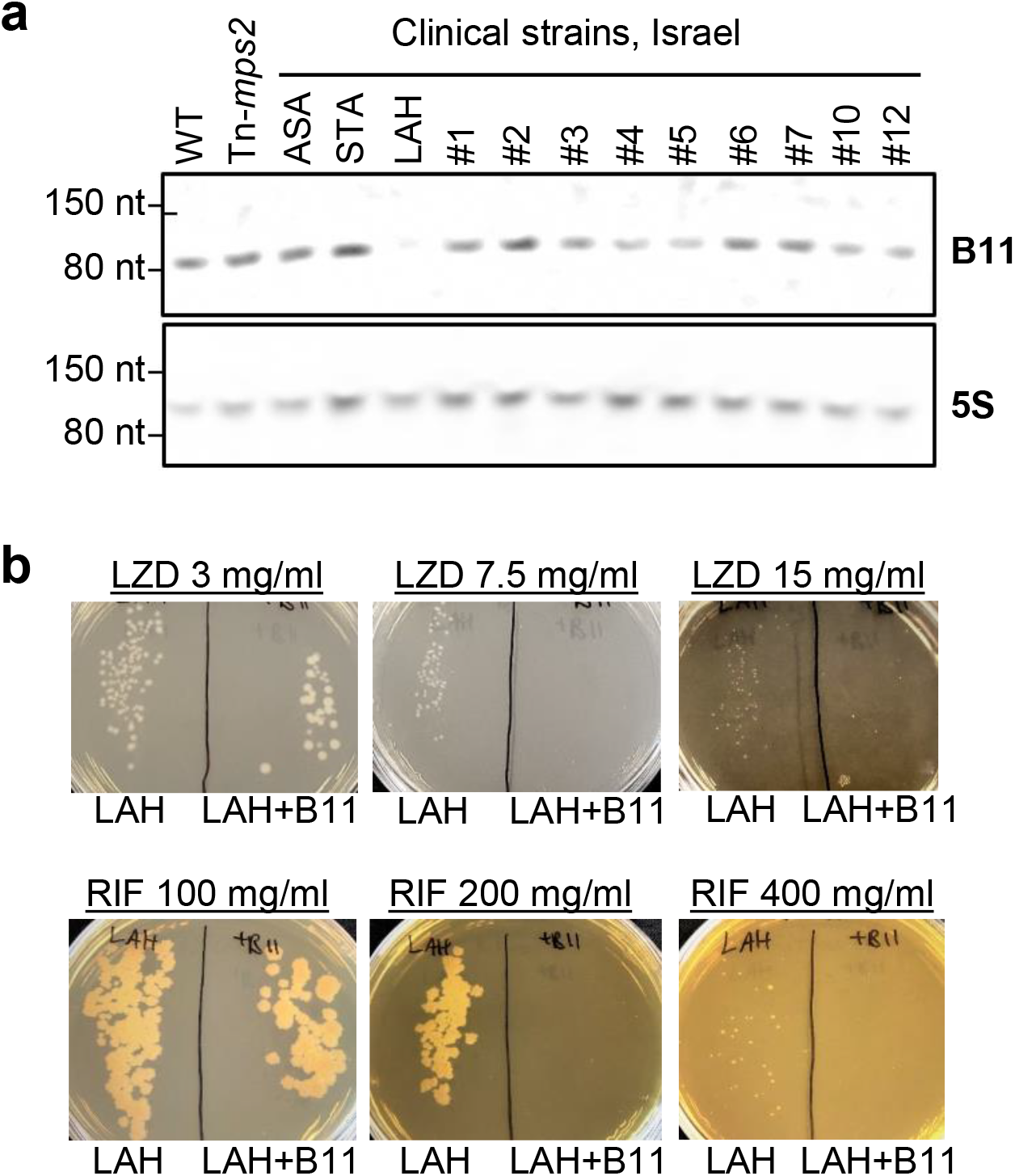
A clinical *M. abscessus* strain does not express B11 and had increased antibiotic sensitivity upon ectopic B11 expression. **A**. Northern blot for B11 in log phase cultures of WT *M. abscessus*, a transposon-mutant of *mps2* (MAB_4098c), and twelve clinical isolates from an Israeli cohort. **B**. The clinical isolate LAH was electroporated with a single copy plasmid encoding B11 under the MOP promoter, and sensitivity to linezolid and rifampin was examined.

## DISCUSSION

In most bacteria, virulence mechanisms (such as ESX-4 in *M. abscessus*) and virulence-related phenotypes (such as the lack of GPL in *M. abscessus*, leading to a rough colony-morphology phenotype) are tightly regulated at multiple levels. Small non-coding RNAs have been shown to play an important role in such regulation in several bacteria, where their role is usually of “fine tuning", rather than overt “all or none” regulation. In mycobacteria, only limited data exist on physiologic roles of sRNA, and even fewer are directly linked to virulence. The mechanisms of genetic regulation by sRNA are also understood to a lesser extent in mycobacteria than in other bacteria, partially due to a lack of identified RNA-chaperone proteins (such as Hfq and ProQ). In this work, we identified that in *M. abscessus* the sRNA B11 affects expression of over 200 genes. B11 appears to negatively regulate some genes directly, while positively regulating others, possibly through indirect mechanisms. Further study is needed to determine which genes are regulated directly at the level of transcript stabilization or destabilization and which are regulated indirectly at the level of transcription. Interestingly, B11 seems to negatively regulate one virulence system (the ESX-4) and positively regulate GPL production (which may be considered an “anti-virulence” system) – collectively making it a suppressor of pathogenesis. Deletion mutants were hyper-virulent in several experimental models (cell-culture, infection of larvae, and a murine model of lung infection).

If B11 is indeed a negative regulator of virulence, a question arises whether B11 downregulation plays a role in the clinical course of human infections. Most patients with CF who eventually develop severe *M. abscessus* lung disease are first colonized by smooth variants. Over time, mutations favoring virulence arise, are naturally selected by the host environment, and accumulate in the *M. abscessus* population. Chronic inflammatory disease ensues – to a great extent due to a destructive immune response of the host. This process entails a different set of genetic events in each clinical isolate of *M. abscessus*. By examining cohorts of over 100 clinical isolates from Ireland, Denmark, and Israel we found one isolate with substantial downregulation of B11 expression, and several isolates with indel mutations in the C-loops of B11 – mutations we show to reduce the biologic activity of the molecule. Taken together, it appears that downregulation of B11 (either through expression or mutations) is one of the genetic mechanisms by which *M. abscessus* evolves *in-vivo* towards a more virulent phenotype. The small but consistent increases in the bacteria’s MIC to linezolid and rifampicin after B11 deletion also stress the role this molecule may play in exacerbating the clinical picture in affected patients.

As we mentioned, the mechanism by which sRNA act in mycobacteria, in the apparent absence of an sRNA chaperone, is poorly understood. In (36), examination of the interaction between the B11 homolog of Mtb with genes of *M. smegmatis* led to the suggestion that direct pairing of bases could occur without need of a chaperone. This work also suggested that B11 may be essential, as a deletion mutant could not be created in *M. smegmatis*. Our results are not in accordance with this hypothesis, as B11 deletion mutants were relatively easily constructed both in *M. abscessus* and in *M. smegmatis* – however our results do agree with the hypothesized direct interaction of either one of the long C-loops with G-rich areas in the UTR of genes regulated by B11 – as both the destruction of the loops or the G-stretches reduced the effect of B11 on reporter genes. Both RNAseq and whole cell proteomic analyses showed genes with long G-stretches in their UTRs are much more likely to be affected by B11 than genes without these G-stretches in the UTRs.

Previously, researchers performing a transposon-mutant screen in *M. kansasii* found a transposon insertion at the equivalent TA position to our Tn-B11 mutant (56). Despite the similarity in the colony-morphology phenotypes, the *M. kansasii* mutant in their study lost the ability to form biofilms, whereas the *M. abscessus* mutant in our study was clumpier, suggesting more potential for biofilm formation. Biofilm formation in rough *M. abscessus* strains has been described previously (57). Budell *et al* also tested pathogenesis in the *G. mellonella* model but found no difference, whereas we found the B11 mutant was hypervirulent. These differences could reflect differences in the roles of B11 in the two species. Alternatively, they could be due to experimental design; Budell *et al* were looking for attenuation and therefore designed the experiment such that the WT control produced maximal virulence, making hypervirulence in a mutant hard to detect. In our study we used a smaller inoculum, allowing the hypervirulent phenotype to be uncovered.

The interplay of B11 with the ESX-4 system is intriguing. ESX-4 secretion was shown some time ago to play an important role in *M. abscessus* pathogenesis (20). Here we showed that B11 deletion causes upregulation of the expression of ESX-4 components, including the structural gene *eccB4*, and the two secretion substrate genes *esxU* and *esxT*. This initially suggested that part of the explanation for the increased virulence could be increased ESX-4 activity. However, a recent paper found that specific deletion of the secretion substrates alone (*esxU* and *esxT*) caused a hypervirulent phenotype in mice, suggesting the role of ESX-4 in virulence is complex and still poorly understood. The exact interplay, therefore, of B11, ESX-4 as a whole, specifically the ESX-4 secretion substrates and bacterial virulence should continue to be a focus of intense research.

## MATERIALS AND METHODS

### Bacteria and growth conditions

*M. abscessus* ATCC_19977 was obtained from the ATCC collection. Clinical isolates were obtained from the Israeli Ministry of Health Mycobacteria Central Laboratories, and from clinical microbiology laboratories in various hospitals. Bacteria were grown in 7H9 media supplemented by 0.05% glycerol, 10% ADC or OADC, and tween 80, as widely described. For solid agar plates, either LB supplemented with 0.05% glycerol and 0.05% dextrose was used, or 7H10 plates with glycerol and ADC or OADC. Antibiotic concentrations were: kanamycin 120-240 μg/ml and zeocin 33-50 μg/ml for *M. abscessus*; kanamycin 40 μg/ml and zeocin 33 μg/ml for *E. coli*.

### Creation of a Tn-mutant library and identification of the Tn-insertion site in selected clones

To construct the transposon, the zeocin and kanamycin resistance genes were cloned next to each other, with the 27 bp of the Himar-1 inverted repeat on either side. The transposase was PCR amplified from MycoMar7 phage (kind gift from Eric Rubin and Chris Sassetti), and cloned next to the transposon, but outside of the IR. This construct was used to create a temperature-sensitive, TM4-based mycobacteriophage, as previously widely described (58). This phage was used to infect *M. abscessus* at 37°C, and bacteria were then plated on 7H10 plates supplemented with zeocin (50 μg/ml) and kanamycin (120 μg/ml). Aberrant-appearing colonies were identified by the naked eye, and analyzed individually. To identify the Tn-insertion site in clones of interest, genomic DNA was extracted, digested by SalI, self-ligated, and a PCR amplifying the genome-IR junction was performed Using primers M6 and M5 (see Table S3). The resulting fragment was sent for Sanger sequencing, identifying the Tn-junction area. Strain names are listed in Table S4.

### RNA extraction, northern blot, and evaluation of gene expression by qPCR

*M. abscessus* log phase cultures were used to inoculate 50 ml conical tubes containing 10 ml of 7H9 medium to an OD_600_ = 0.01. Cultures were grown at 37°C and 200 RPM until they reached an OD_600_ between 0.6-0.8 and were frozen with liquid nitrogen and stored at −80°C until RNA purification. RNA was extracted as in (59). Briefly, frozen cultures were thawed on ice and centrifuged at 4,000 rpm for 5 min at 4°C. The pellets were resuspended in 1 ml Trizol (Life Technologies) and placed in tubes containing Lysing Matrix B (MP Bio). Cells were lysed by bead-beating twice for 40 s at 9 m/sec in a FastPrep 5G instrument (MP Bio). RNA was purified using Direct Zol RNA miniprep kit (Zymo) according to manufacturer’s instructions. RNA concentrations were determined using a Nanodrop instrument. RNA samples were stored at −80°C until use.

The relative abundance of B11 transcript among strains was evaluated by northern blot as follows: 5 μg of each RNA sample (or 1 μg for 5S, used as a load control) were mixed with TBE-Urea sample buffer (Novex) and run in a 10% TBE-Urea Polyacrylamide gel (Biorad) at 180 V for 1 hour. RNA was transferred to a positively charged nylon membrane (Amhersham) during 80 min at 10 V and 400 mA and then RNA was cross-linked by exposure to UV light (302 nm) for 7 min. Prior to probe hybridization, the membrane was incubated with 10 ml of ULTRAhyb buffer (Ambion) at 50 °C for 30 min. Then, the membrane was incubated with 10 ml of ULTRAhyb buffer containing ~ 200 ng of B11 RNA-probe or 5S RNA-probe (see Table S3) at 50°C overnight. The membrane was washed with 30 ml of low stringency wash solution (2X SSC, 0.1% SDS) for 10 min at room temperature, then washed with 30 ml of high stringency wash solution (0.1 SSC, 0.1% SDS) at 50°C for 15 min and finally washed with 20 ml washing buffer (Roche) at room temperature for 5 min. The membrane was incubated with 20 ml of blocking solution (Roche) for 30 min and then incubated with 20 ml of antibody solution (Roche) containing 1 μL of Anti-digoxigenin-AP conjugate (Roche) at room temperature for 30 min. The membrane was washed twice with 20 ml of washing buffer and then incubated with 20 ml of detection buffer for 3 min. Detection was done by incubation with 1 ml of detection buffer containing 100 μL CDP-Star (Roche) and exposure in a Gel Doc.

Expression of *MAB_3759c*, *MAB_3753c*, *MAB_3754c*, *MAB_0498c*, *MAB_0499c, MAB_0628*, and *MAB_4938* relative to *sigA* was determined by qPCR using the primers listed in Table S3. cDNAs were prepared as described (59). For each qPCR reaction, iTaq SYBR green (Bio-Rad) was mixed with 200 pg cDNA and 0.25 μM each primer in 10 μl reaction mixtures. The qPCR parameters were: 40 cycles of 15 s at 95°C and 1 min at 61°C.

### RNAseq

rRNA was depleted and paired-end Illumina sequencing libraries were constructed as described (60,61). Libraries were sequenced at the UMass Medical School Deep Sequencing Core Facility on a HiSeq 2000. Raw fastq files were demultiplexed using Cutadapt (62) by providing corresponding sample barcodes. Reads were mapped to the NC_010397 reference genome using Burrows-Wheeler Aligner mem (63). The FeatureCounts tool was used to assign mapped reads to genomic features for producing count matrices (64). Principal Component Analysis (PCA) was applied for quality assessment of RNA expression libraries. Deseq2 (65) was used to assess changes in gene expression. The Deseq2 package internally corrects for library size. The input count matrix for differential expression analysis were un-normalized counts, which allow assessing the measurement precision correctly. Raw and processed data are available in GEO, accession number GSE214640.

### Targeted B11 deletion in *M. abscessus*

*M. abscessus* ATCC19977 was first electroporated with the plasmid pJV53 (kanamycin), with recombineering enzymes under an acetamide-regulated promoter (66), to enhance targeted deletion events when using a specialized TM4-based mycobacteriophage (67). The 5’ and 3’ flanking regions (600 bp long) of B11 were cloned on either side of the zeocin^R^ gene, to create pDB386. This plasmid was used to create the specialized transducing phage phDB35 (58). *M. abscessus*^pJV53^ was then grown in media with 0.2% succinate and no dextrose, induced by acetamide for 4 hours, and infected by phDB35 for 3 hours at 37°C. Bacteria were then washed, re-suspended in 7H9 media, and plated on 7H10 plates with 50 μg/ml zeocin. One of multiple zeocin-resistant colonies was further examined by PCR reactions with the primers yielding a 1500 bp product in WT and a 1900 bp product in a mutant where a correct replacement took place (Fig. S1c) Sanger sequencing of the product was also performed. After confirmation of the deletion, the mutant was grown and plated for single colonies. Colonies were then patched with and without kanamycin, to verify the loss of pJV53. A kanamycin-sensitive colony was isolated, and the final strain, *M. abscessus* ΔB11, was named mDB228.

### Complementation of the B11 deletion in *M. abscessus*

The region spanning from 250 bp upstream to 200 bp downstream of B11 was PCR-amplified and cloned into the L5 *attB*-integrating, kanamycin-selected plasmid pDB213, to create pDB392. This plasmid was used to complement the ΔB11 mutant (mDB228), to create a complemented mutant mDB252 (ΔB11^zeo^+B11^kan^). PCR-mutagenesis was used to introduce a C deletion and a C insertion in the first C-rich loop and second C-rich loop of B11, respectively, in pDB213 (Fig.5b), and the resulting plasmids were transformed into mDB228, to create the complemented mutants mDB281 (ΔB11^zeo^+B11_del1_^kan^) and mDB282 (ΔB11^zeo^+B11_del1_^kan^). PCR-mutagenesis was also used to introduce multiple base substitutions into the second C-loop of B11 (Fig. S1e), to create mDB269 (ΔB11^zeo^+B11_mutated_^kan^).

### Construction of the reporter systems

For the reporter system described in Figure 5, *M. abscessus* ATCC19977 and the ΔB11 mutant were electroporated with a set of reporters where mCherry was expressed under the MOP promoter either with the 55 nt that correspond to the MOP-associated synthetic 5’ UTR (P_MOP__UTR_MOP__mCherry) or with the 74 nt that correspond to the eccB4 5’ UTR (P_MOP__UTR_eccB4__mCherry) (Fig 5b-c). Then, to evaluate which loop of B11 was responsible for its mediated regulation, different versions of B11 were cloned into the P_MOP__UTR_eccB4__mCherry construct divergently from P_MOP_. PCR was used to amplify B11 and its native promoter (223 nt upstream B11) and the obtained fragment was cloned into the P_MOP__UTR_eccB4__mCherry reporter system to generate the B11_P_B11__B11_P_MOP__UTR_eccB4__mCherry construct. PCR-mutagenesis was later used to introduce multiple versions of base substitutions into the first, second or both loops, as described in the text (Fig 5b-d). All constructs were done on kanamycin-selected L5-integrating plasmids.

For the reporter system described in Figure 6, we first cloned a B11-neutral, similar mCherry construct as described above (P_MOP__UTR_MOP__mCherry) into the HpaI site of pDB213 (a kanamycin-selected, L5 integrating plasmid), to create pDB266. We then cloned mWasabi with the 74 nt UTR of *eccB4* (P_MOP__UTR_eccB4__mWasabi) into the AvrII site of pDB266 (version α in the figure). Of note, this also disrupted the *integrase* gene in pDB266, making the integration into the *attB* site more difficult and dependent on supplementation of the integrase on another plasmid, but also much more stable once completed. PCR-mutagenesis was later used to introduce three point mutations in the six guanosines rights before the ATG start codon, changing them into 5’-GTGTGT-3’ (version β in the figure).

### Flow cytometry

Three biological triplicates of each strain were used to evaluate mCherry and/or mWasabi expression by flow cytometry. Cultures were grown in 7H9 media with glycerol to an OD_600_ of 0.5, 2 ml of cultures were centrifuged, and the bacterial pellets were resuspended in 4% paraformaldehyde (Thermo Scientific). After 1 h of incubation at room temperature cultures were washed twice with PBST (PBS + 0.1% Tween 20, freshly filtered with a 0.2 μm filter to remove precipitates that otherwise appear as bacteria-like events on the flow cytometer) and resuspended in 2 mL of freshly filtered 7H9 media with glycerol. Cultures were filtered using 5 μm filter needles (BD Nokor filter needle) to remove clumps and a final dilution using freshly filtered 7H9 media with glycerol was done to achieve a final concentration of OD=0.025. Samples were analyzed with a Cytoflex S (Beckman Counter) or NovoCyte Quanteon (Agilent) flow cytometer. The parameter settings were for Gain: FSC 500, Violet SSC 50 and for Thresholds: Violet SSC-H 100,000, FSC-H 10,000. Filtered 7H9 media was used to wash between samples and 5000 events were recorded within a gate selected to capture individual cells. Data were analyzed using the FlowJo V10 Software and the median of the detected fluorescence was compared between strains.

### Proteomic analysis of *M. abscessus* mutants

For proteomic analysis of bacteria, *M. abscessus* was grown in 7H9 media with glycerol but no albumin to an O.D600 of 0.3, 10 ml of cultures were centrifuged, and the bacterial pellet was stored at −80°C. Sample preparation and LC-MS/MS were performed at the Smolar Centre for Proteomics, Technion, Haifa, Israel as follows: The samples were mixed with 8 M Urea, 100 mM ABC, and 2.8 mM DTT (60°C for 30 min), modified with 8.8 mM iodoacetamide in 100 mM ammonium bicarbonate (in the dark, room temperature for 30 min) and digested in 2 M Urea, 25 mM ammonium bicarbonate with modified trypsin (Promega) at a 1:50 enzyme-to-substrate ratio, overnight at 37°C. An additional second digestion was done for 4 hours. The resulting peptides were desalted using C18 tips (Homemade stage tips) dried and re-suspended in 0.1% Formic acid. The peptides were resolved by reverse-phase chromatography on 0.075 × 180-mm fused silica capillaries (J&W) packed with Reprosil reversed phase material (Dr Maisch GmbH, Germany). The peptides were eluted with a linear 60 minute gradient of 5 to 28%, 15 minutes gradient of 28 to 95%, and 15 minutes at 95% acetonitrile with 0.1% formic acid in water at flow rates of 0.15 μl/min. Mass spectrometry was performed by Q Exactive plus mass spectrometer (Thermo) in a positive mode using repetitively full MS scan followed by collision induces dissociation (HCD) of the 10 most dominant ions selected from the first MS scan.

The mass spectrometry data were analyzed using the Protein Discoverer 1.4 (Thermo) using Sequest search engine, searching against *M. abscessus* of the Uniprot database. Peptide- and protein-level false discovery rates (FDRs) were filtered to 1% using the target-decoy strategy.

### Protein secretion assay

*M. abscessus* ATCC_19977, ΔB11 mutant, and ΔB11+B11 were grown to an OD_600_ of 0.8 in 7H9 media with glycerol, washed twice with PBS, resuspended in Sauton media (KH_2_PO_4_ 0.5 g, MgSO_4_.7H_2_O 0.5 g, citric acid 2.0 g, ferric ammonium citrate 0.05 g, glycerol 60 ml, asparagine 4.0 g per L) supplemented with 0.05% Tween 80 and grown for 24 hours to an OD_600_ of ~0.6-0.8. Cultures were pelleted, and whole cell lysates were made by resuspending the pellets in 500 μl protein extraction buffer (50 mM Tris pH 7.5, 5 mM EDTA, and 1X protease inhibitor cocktail with 0.5 M EDTA), transferring to lysing matrix B tubes (MP Bio), and disrupting by bead-beating in a Fastprep 5G instrument (MP Bio) for 4 cycles of 40 seconds at 9 m/sec and 2 min incubations on ice between cycles. After cell disruption, 170 μl of SDS-PAGE loading buffer (200 mM Tris pH 6.8, 400 mM DTT, 8% SDS, 0.4% bromophenol blue, 40% glycerol) was added to whole cell lysates.

In parallel, culture supernatants were combined with 100 μl protease inhibitor cocktail (Thermo Scientific) with 0.5 M EDTA, filtered through 0.2 μm pore size filters (Genesee Scientific), and incubated overnight with 5 mL of concentrated trichloroacetic acid (Chemicals BDH) at 4°C. Then proteins were pelleted by 20 minutes of centrifugation at 14,500 RPM at 4°C, washed once with 100% acetone (Chemicals BDH), centrifuged, and finally resuspended in 100 μl protein extraction buffer plus 100 μl SDS-PAGE loading buffer. Both whole cell lysate and culture supernatant samples were heated at 95°C for 5 minutes before running on 4-20% PAGE gels in tris-glycine-SDS buffer and staining with Coomassie Blue.

Protein secretion assays were also performed using *M. abscessus* ATCC19977, ΔB11 mutant, and tn-*mps2* transformed with a plasmid expressing mCherry (P_MOP__UTR_MOP__mCherry). Whole cell lysate and supernatant of two biological replicates were probed by western blotting using a 1:1000 dilution anti-mCherry antibody (Proteintech) as a primary antibody and a 1:30000 dilution goat anti-rabbit antibody as a secondary antibody. Detection was performed by incubation with Radiance ECL (Azure biosystems) according to manufacturer’s instructions and exposure in a Gel Doc.

### Infection of RAW cells and TNF secretion measurement

RAW 264.7 macrophages were cultured in DMEM (ATCC) supplemented with 10% fetal bovine serum. Prior to infection, RAW 264.7 cells were seeded in 6-well plates (2 × 10^5^ per well). For experiments on exposure to heat-killed bacteria, heat-killed (30 minutes in 65°C) *M. abscessus* ATCC_19977 and ΔB11 mutant were prepared. RAW 264.7 were exposed to the bacteria at a multiplicity of infection (MOI) of 20. After incubation for 4 hours, cells were washed with phosphate-buffered saline (PBS) three times and fresh culture medium was added. Following 24 hours of incubation, culture medium was collected and analyzed for murine TNF by ELISA kit (Invitrogen, 88-7324) according to manufacturer’s instructions. For live-bacteria infection experiments, RAW cells were prepared and seeded the same way. Live *M. abscessus* ATCC_19977 and ΔB11 mutant were prepared and used to infect the RAW cells at an MOI of 5. After an incubation of 6 hours, the culture medium was collected for TNF measurement by a similar ELISA kit.

### Murine infections

Animal studies were conducted according to protocols and adhering strictly to the Italian Ministry of Health guidelines for the use and care of experimental animals (IACUC N°816) and approved by the San Raffaele Scientific Institute Institutional Animal Care and Use Committee (IACUC). Immunocompetent C57BL/6NCrl male mice (8 to 10 weeks of age) from Charles River were used for all experiments. Mice were maintained in specific pathogen-free conditions at San Raffaele Scientific Institute, Milan, Italy. *M. abscessus* ATCC_19977 and the ΔB11 mutant were streaked on 7H10 plates and colonies used to inoculate 20 ml of Middlebrook 7H9 broth. After two days of growth, the bacteria were embedded in agar beads as previously described (44,45,68), mice were anesthetized, the trachea was exposed and intubated, and 50 μL of beads suspension (10^5^ CFU) were injected before closing the incision with suture clips. Control mice were intratracheally inoculated with the same volume of empty beads suspension. After infection, mice were monitored daily for body weight, appetite and hair coat. Four days after infection, mice were euthanized by CO_2_ asphyxiation. Lung and spleen were collected, homogenized, and processed for microbiological analysis. Total CFU were the result of the addition of the CFU in lung homogenate and bronchoalveolar lavage fluid. Lung homogenates were centrifuged at 16,000 × g for 30 min at 4°C, and then stored at −80°C for cytokines and chemokines analysis. TNF, KC (CXCL1), and MIP-2 (CXCL2) concentrations were evaluated in the lung homogenate supernatants by DuoSet ELISA Development Systems (R&D Systems), according to the manufacturer’s instructions.

### Infection of *Galleria mellonella*

The *G. mellonella* model was used as previously described (42). Briefly, larvae were infected by a 30G needle, with 500-1000 CFU of *M. abscessus*, and kept at 30°C. Mortality was monitored daily. Where CFU count was needed, larvae were killed by freezing for 5 minutes at −20°C, de-contaminated in 70% ethanol, homogenized in 5 ml of PBS, and plated in dilutions on appropriate agar plates.

### Determination of MICs for ATCC_19977, ΔB11 and ΔB11+B11 strains, the clinical isolate LAH, and derivative and LAH+B11^MOP^

A luminescent mutant of ATCC_19977 was already created and described by us (42,69). We created a similar luminescent mutant on a ΔB11 background, using the same integrating, kanamycin-resistance vector (pLux13). By cloning B11 with the native promoter into pLux13, we created a ΔB11+B11, luminescent mutant. 500 CFU of each were inoculated into 200 μl of 7H9 media, with serial dilutions of the designated antibiotics, in a luminescence-compatible 96-well plate. Bacteria were kept at 37°C, and growth was monitored daily by measurement of the luminescence (Spectramax © plate reader).

To complement the clinical strain LAH by B11, we first cloned B11 under the constitutive promoter MOP, so it is not dependent on the physiologic regulation of B11 in LAH, that appeared to be downregulated. MOP-B11 was then cloned into an integrating vector (pDB406), selected by both zeocin and kanamycin, to ensure selection in the highly resistant parent strain. pDB406 was electroporated into LAH to create LAH+B11^MOP^. For lack of additional effective selection markers, this mutant could not be made luminescent. To test for MIC, we therefore plated LAH and LAH+B11^MOP^ on LB/dextrose/glycerol plates supplemented by the designated concentration of antibiotics.

### Identification of mutations in clinical isolates

The genomes reported in (54) were analyzed. Fastq files were aligned to NC_010397.1 with bwa (63) and sequence variants identified by Samtools and Bamtools (70). Read depth was verified with Artemis (71). Additionally, genomes of isolates from Denmark were analyzed using a similar approach.

## AUTHOR CONTRIBUTIONS

M.B.O., M.C.M, M.N.A., S.S.S., D.M.C., H.K.J and D.B. conceived and designed experiments. M.B.O., M.C.M., M.N.A., C.R., N.I.L, P.M, and M.M performed experiments. J.X., M.A.M, C.S.M., H.S., and J.K.M. performed bioinformatics analyses. All authors analyzed and interpreted data. M.B.O., M.C.M, M.M, S.S.S., and D.B. wrote the manuscript.

## ACKNOWLEDGEMENTS

We thank members of the Shell and Barkan labs, M. Chiacchiaretta (IRCCS San Raffaele Scientific Institute) and F. Cugnata (Vita-Salute San Raffaele University, Milano, Italy) for technical assistance and helpful discussions. We thank Christina Stallings for critical reading of the manuscript, Margaret Fitzgibbon (St. James Hospital, Dublin) for providing information on the Irish cohort of *M. abscessus*, and Efrat Rorman (Israeli Ministry of Health) for providing some of the clinical isolates in the Israeli cohort.

## Supplementary tables

**Table S1. Results from DESeq2 differential expression analysis, and identification of B11-complementary sequences in the ribosome-binding regions of genes with defined 5’ UTRs**

**Table S2. Results from DESeq2 differential expression analysis and identification of B11-complementary sequences in the ribosome-binding regions of genes with defined 5’ UTRs, for GPL biosynthesis and transport genes only.**

**Table S3. Sequences of primers and northern blot probes used in this work.**

**Table S4. Strains used in this work.**

---

**Figure S1.**
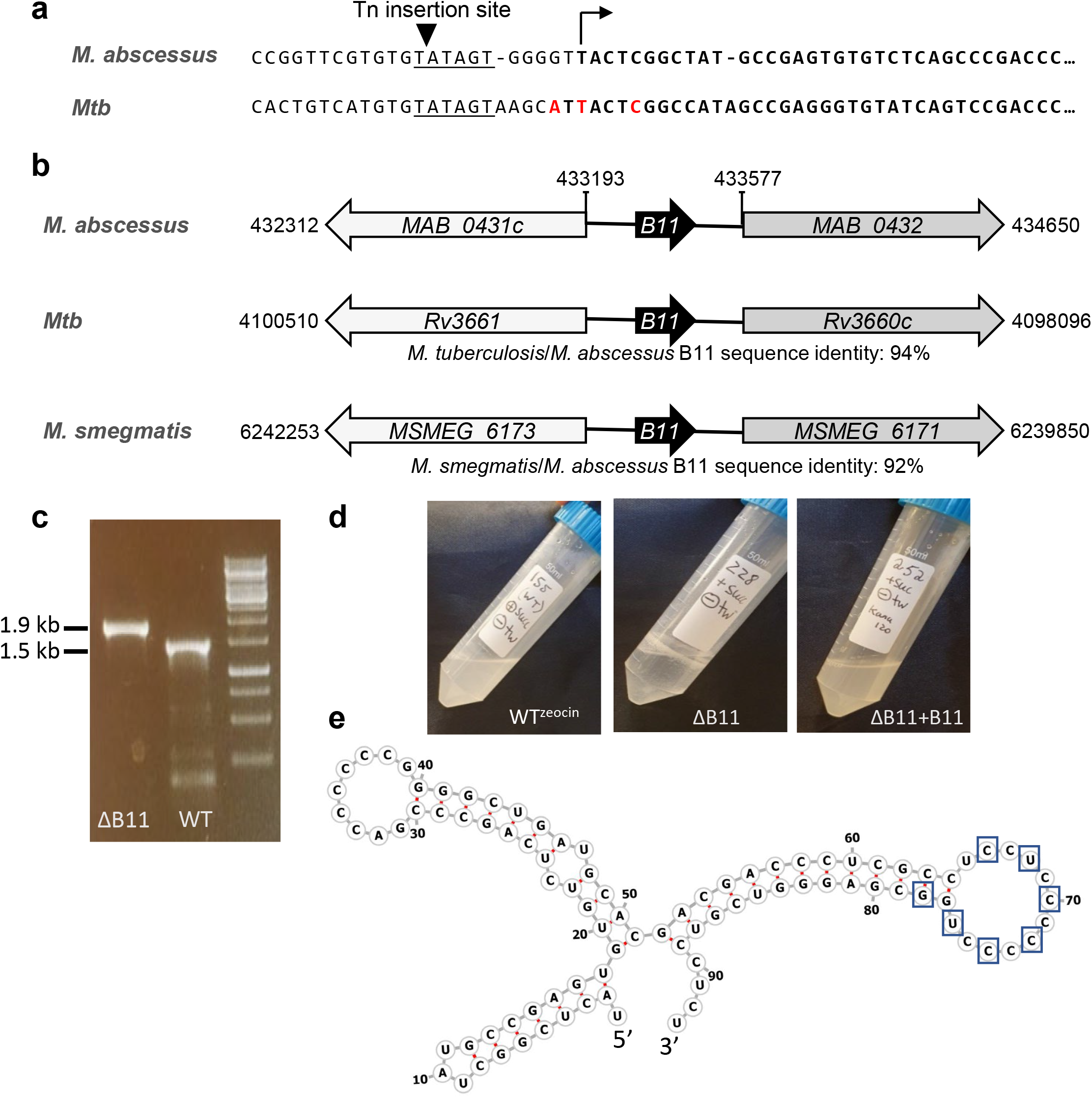
Disruption of the sRNA B11 by a transposon and by targeted replacement. **A**. Schematic showing the position of transposon insertion at the *M. abscessus* B11 locus, and alignment of the surrounding sequence with the equivalent portion of the *M. tuberculosis* B11 locus. **B**. Genomic context of B11 in three mycobacterial species. Not to scale. **C**. PCR reactions with primers binding 700 bp upstream and downstream of B11 were performed on WT and on the deletion candidate. The reaction yields a 1500 bp product in WT, and a 1900 bp product when B11 is replaced by the 500 bp zeocin^R^ gene. **D**. Deletion of B11 caused extensive clumping during growth in liquid media without Tween. **E**. Predicted secondary structure of B11 (Vienna RNAfold). Boxed positions were mutated as follows in B11_mutated_ shown in Fig 1a and 4b: G to A, C to U, and U to C.

**Figure S2.**
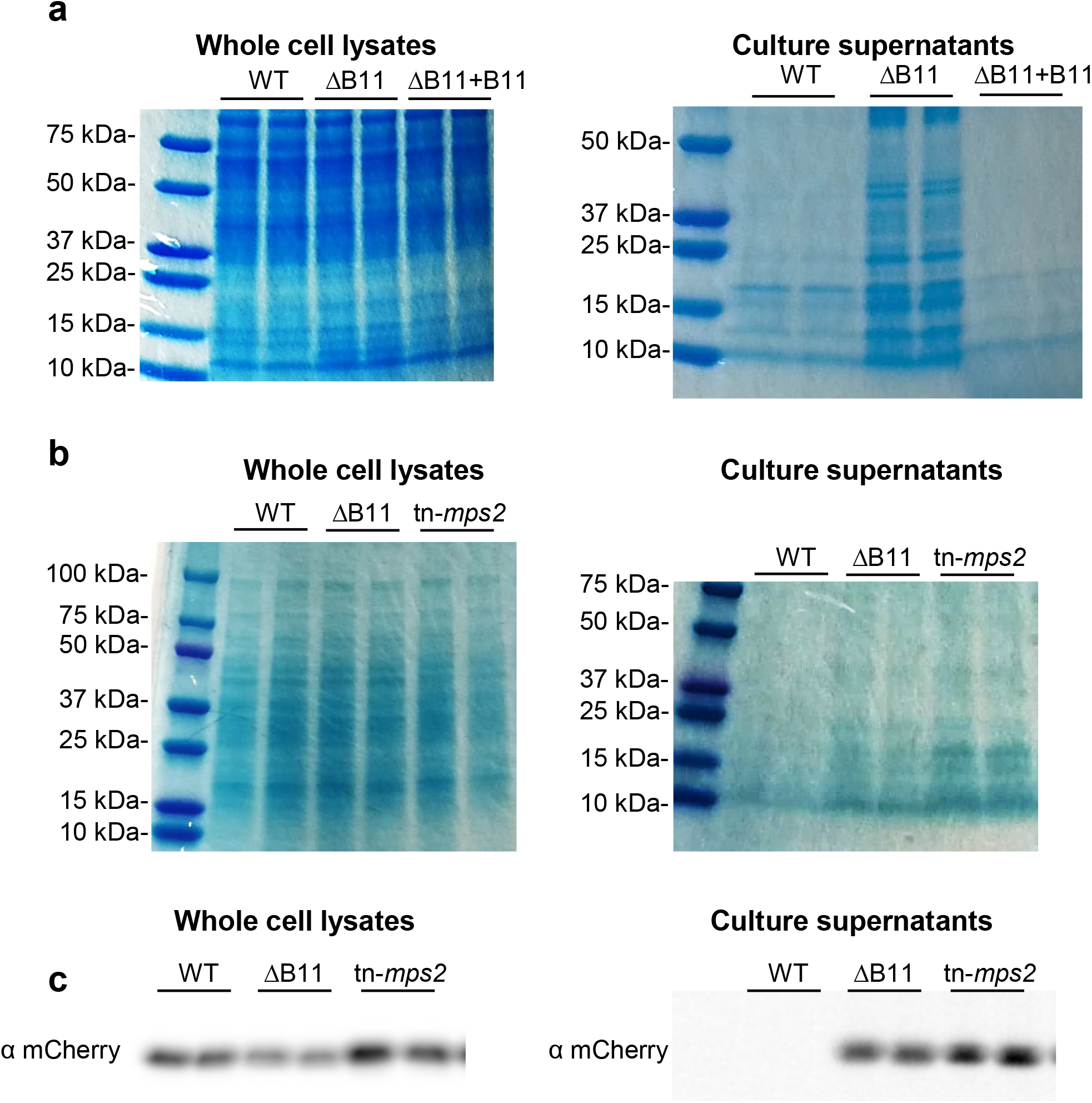
A B11 deletion strain and a GPL biosynthesis mutant both release more protein into culture supernatants than smooth strains. **A**. Protein was extracted from cell pellets or from culture supernatants. Quantities from equivalent culture volumes were separated by SDS-PAGE and stained with Coomassie blue. The ΔB11 strain consistently released more protein into the culture supernatant than the WT or complemented strains. **B**. The same assay revealed that a rough strain with a transposon disruption of *msp2* also released more protein into culture supernatants than the smooth WT parent. **C**. The indicated strains were transformed with a plasmid expressing mCherry (which lacks a secretion signal sequence) and their cell lysates and culture supernatants were probed by western blotting against mCherry. mCherry was exclusively cytoplasmic in the WT strain as expected, but present at detectable levels in the culture supernatants of the rough strains.

**Figure S3.**
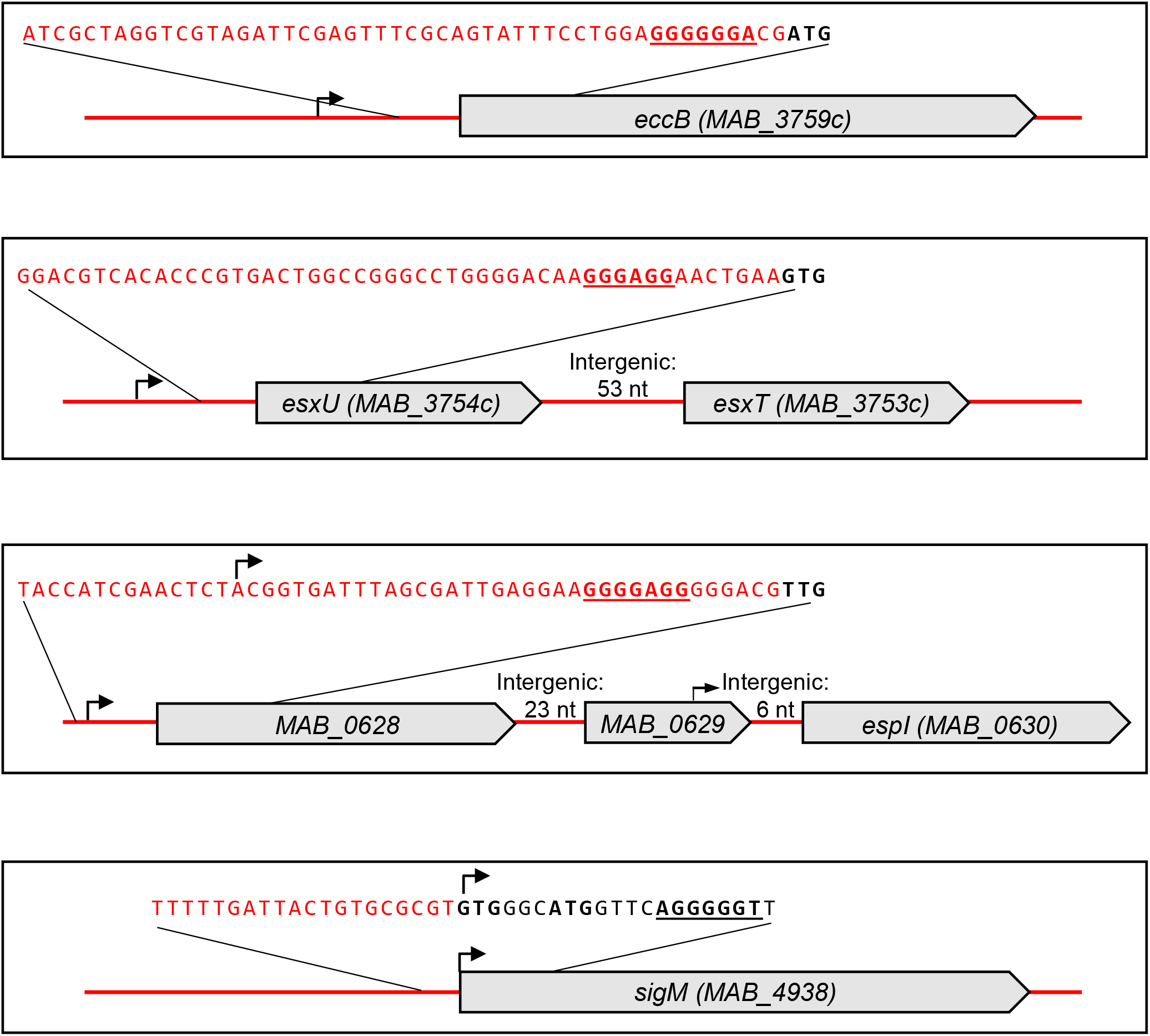
The genetic architecture of selected ESX-related genes repressed by B11. Coding sequences are shown in black and intergenic sequences are shown in red The 50 nt sequence upstream of each start codon is shown. Start codons are bolded Regions 6 nt or longer of complementarity to either loop of B11 are bolded and underlined. Shine-Dalgarno sequences in mycobacteria are typically located between −13 and −7 relative to the start codon (Shell et al 2015, Martini et al 2019, and our unpublished analyses). Published transcription start sites (Miranda-CasoLuengo et al 2016) are indicated with bent arrows. Note that the TSS proximal to *MAB_0630* had much lower read depth compared to the TSS upstream of *MAB_0628*, suggesting that the genes may be transcribed primarily as a polycistron. The *sigM* gene is likely transcribed as a leaderless mRNA; its annotated 5’ UTR is 6 nt, but the transcript begins with an in-frame GTG and we have previously shown that such transcripts are translated from their 5’ ends (Shell et al 2015). Both the annotated (downstream) and corrected (upstream) star codons are shown here. A sequence in the coding region that is complementary to 6 nt o B11 loop 1 or 6 nt of B11 loop 2 is bolded and underlined. Elements are not drawn to scale.

**Figure S4.**
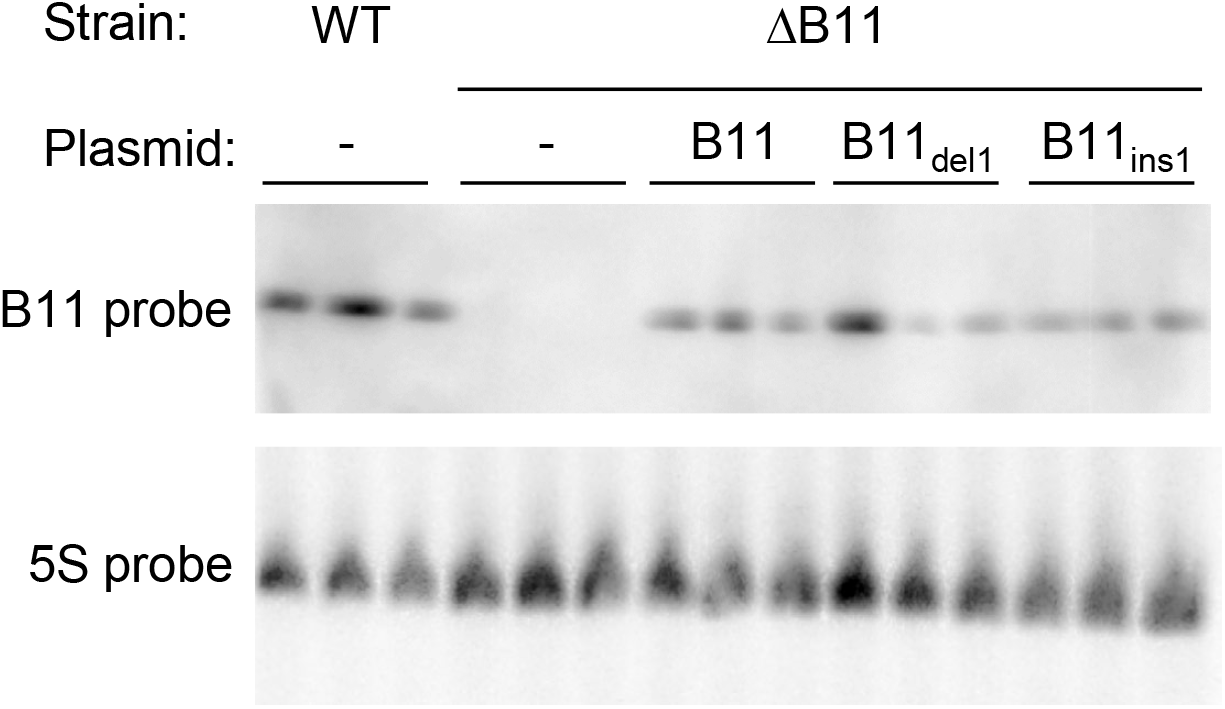
B11 mutations found in clinical *M. abscessus* strains do not affect abundance when expressed ectopically in a B11 deletion strain. Total RNA from triplicate cultures of the indicated strains was transferred to a membrane and probed sequentially for B11 and 5S rRNA as a loading control.

**Figure S5.**
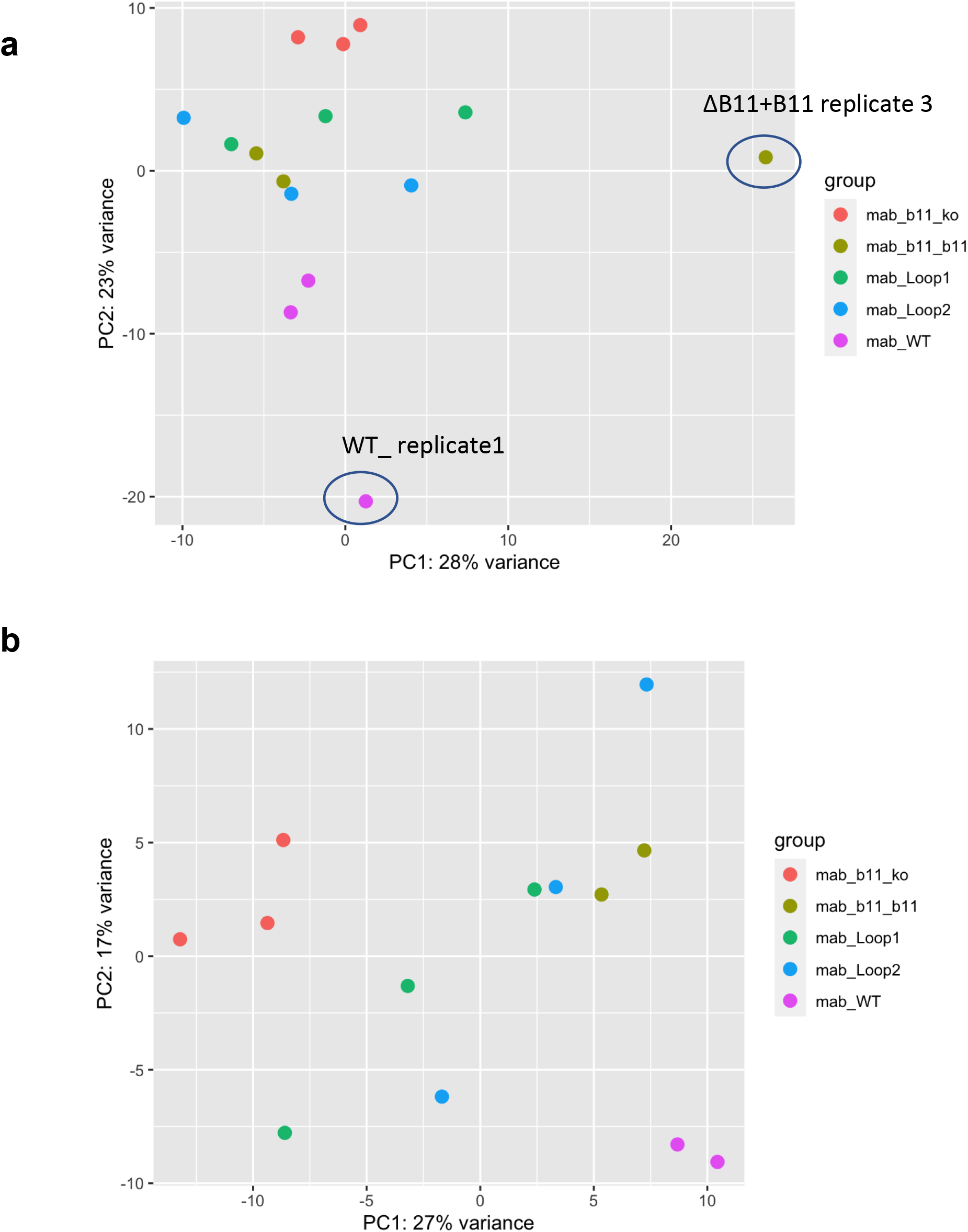
PCA of RNAseq read counts. **A**. All three replicates of the five indicated strain were clustered based on read counts generated by FeatureCount. The circled replicate were eliminated from further analysis due to being outliers. Additionally, the eliminate WT replicate had a very low read count and the eliminated complemented replicat lacked detectable expression of B11. **B**. The remaining samples were clustered afte removed of the samples indicated in **A**.

## REFERENCES

1. Maiz, L., Giron, R., Olveira, C., Vendrell, M., Nieto, R., and Martinez-Garcia, M.A. (2016) Prevalence and factors associated with nontuberculous mycobacteria in non-cystic fibrosis bronchiectasis: a multicenter observational study. BMC Infect Dis, 16, 437.

2. Olivier, K.N., Weber, D.J., Wallace, R.J., Jr., Faiz, A.R., Lee, J.H., Zhang, Y., Brown-Elliot, B.A., Handler, A., Wilson, R.W., Schechter, M.S., et al. (2003) Nontuberculous mycobacteria. I: multicenter prevalence study in cystic fibrosis. Am J Respir Crit Care Med, 167, 828–834.

3. Roux, A.L., Catherinot, E., Ripoll, F., Soismier, N., Macheras, E., Ravilly, S., Bellis, G., Vibet, M.A., Le Roux, E., Lemonnier, L., et al. (2009) Multicenter study of prevalence of nontuberculous mycobacteria in patients with cystic fibrosis in france. J Clin Microbiol, 47, 4124–4128.

4. Novosad, S.A., Beekmann, S.E., Polgreen, P.M., Mackey, K., Winthrop, K.L., and Team, M.a.S. (2016) Treatment of Mycobacterium abscessus Infection. Emerg Infect Dis, 22, 511–514.

5. Floto, R.A., Olivier, K.N., Saiman, L., Daley, C.L., Herrmann, J.L., Nick, J.A., Noone, P.G., Bilton, D., Corris, P., Gibson, R.L., et al. (2016) US Cystic Fibrosis Foundation and European Cystic Fibrosis Society consensus recommendations for the management of non-tuberculous mycobacteria in individuals with cystic fibrosis: executive summary. Thorax, 71, 88–90.

6. Furukawa, B.S., and Flume, P.A. (2018) Nontuberculous Mycobacteria in Cystic Fibrosis. Semin Respir Crit Care Med, 39, 383–391.

7. Qvist, T., Taylor-Robinson, D., Waldmann, E., Olesen, H.V., Hansen, C.R., Mathiesen, I.H., Hoiby, N., Katzenstein, T.L., Smyth, R.L., Diggle, P.J., et al. (2016) Comparing the harmful effects of nontuberculous mycobacteria and Gram negative bacteria on lung function in patients with cystic fibrosis. J Cyst Fibros, 15, 380–385.

8. Park, I.K., Hsu, A.P., Tettelin, H., Shallom, S.J., Drake, S.K., Ding, L., Wu, U.I., Adamo, N., Prevots, D.R., Olivier, K.N., et al. (2015) Clonal Diversification and Changes in Lipid Traits and Colony Morphology in Mycobacterium abscessus Clinical Isolates. J Clin Microbiol, 53, 3438–3447.

9. Kreutzfeldt, K.M., McAdam, P.R., Claxton, P., Holmes, A., Seagar, A.L., Laurenson, I.F., and Fitzgerald, J.R. (2013) Molecular longitudinal tracking of Mycobacterium abscessus spp. during chronic infection of the human lung. PLoS One, 8, e63237.

10. Roux, A.L., Viljoen, A., Bah, A., Simeone, R., Bernut, A., Laencina, L., Deramaudt, T., Rottman, M., Gaillard, J.L., Majlessi, L., et al. (2016) The distinct fate of smooth and rough Mycobacterium abscessus variants inside macrophages. Open Biol, 6.

11. Nessar, R., Reyrat, J.M., Davidson, L.B., and Byrd, T.F. (2011) Deletion of the mmpL4b gene in the Mycobacterium abscessus glycopeptidolipid biosynthetic pathway results in loss of surface colonization capability, but enhanced ability to replicate in human macrophages and stimulate their innate immune response. Microbiology (Reading), 157, 1187–1195.

12. Nishimura, T., Shimoda, M., Tamizu, E., Uno, S., Uwamino, Y., Kashimura, S., Yano, I., and Hasegawa, N. (2020) The rough colony morphotype of Mycobacterium avium exhibits high virulence in human macrophages and mice. J Med Microbiol, 69, 1020–1033.

13. Davidson, L.B., Nessar, R., Kempaiah, P., Perkins, D.J., and Byrd, T.F. (2011) Mycobacterium abscessus glycopeptidolipid prevents respiratory epithelial TLR2 signaling as measured by HbetaD2 gene expression and IL-8 release. PLoS One, 6, e29148.

14. Rhoades, E.R., Archambault, A.S., Greendyke, R., Hsu, F.F., Streeter, C., and Byrd, T.F. (2009) Mycobacterium abscessus Glycopeptidolipids mask underlying cell wall phosphatidyl-myo-inositol mannosides blocking induction of human macrophage TNF-alpha by preventing interaction with TLR2. J Immunol, 183, 1997–2007.

15. Bernut, A., Herrmann, J.L., Kissa, K., Dubremetz, J.F., Gaillard, J.L., Lutfalla, G., and Kremer, L. (2014) Mycobacterium abscessus cording prevents phagocytosis and promotes abscess formation. Proc Natl Acad Sci U S A, 111, E943–952.

16. Howard, S.T., Rhoades, E., Recht, J., Pang, X., Alsup, A., Kolter, R., Lyons, C.R., and Byrd, T.F. (2006) Spontaneous reversion of Mycobacterium abscessus from a smooth to a rough morphotype is associated with reduced expression of glycopeptidolipid and reacquisition of an invasive phenotype. Microbiology (Reading), 152, 1581–1590.

17. Catherinot, E., Clarissou, J., Etienne, G., Ripoll, F., Emile, J.F., Daffe, M., Perronne, C., Soudais, C., Gaillard, J.L., and Rottman, M. (2007) Hypervirulence of a rough variant of the Mycobacterium abscessus type strain. Infect Immun, 75, 1055–1058.

18. Pawlik, A., Garnier, G., Orgeur, M., Tong, P., Lohan, A., Le Chevalier, F., Sapriel, G., Roux, A.L., Conlon, K., Honore, N., et al. (2013) Identification and characterization of the genetic changes responsible for the characteristic smooth-to-rough morphotype alterations of clinically persistent Mycobacterium abscessus. Mol Microbiol, 90, 612–629.

19. Steindor, M., Nkwouano, V., Stefanski, A., Stuehler, K., Ioerger, T.R., Bogumil, D., Jacobsen, M., Mackenzie, C.R., and Kalscheuer, R. (2019) A proteomics approach for the identification of species-specific immunogenic proteins in the Mycobacterium abscessus complex. Microbes Infect, 21, 154–162.

20. Laencina, L., Dubois, V., Le Moigne, V., Viljoen, A., Majlessi, L., Pritchard, J., Bernut, A., Piel, L., Roux, A.L., Gaillard, J.L., et al. (2018) Identification of genes required for Mycobacterium abscessus growth in vivo with a prominent role of the ESX-4 locus. Proc Natl Acad Sci U S A, 115, E1002–E1011.

21. Ross, J.A., Thorsing, M., Lillebaek, E.M.S., Teixeira Dos Santos, P., and Kallipolitis, B.H. (2019) The LhrC sRNAs control expression of T cell-stimulating antigen TcsA in Listeria monocytogenes by decreasing tcsA mRNA stability. RNA Biol, 16, 270–281.

22. Lalaouna, D., Baude, J., Wu, Z., Tomasini, A., Chicher, J., Marzi, S., Vandenesch, F., Romby, P., Caldelari, I., and Moreau, K. (2019) RsaC sRNA modulates the oxidative stress response of Staphylococcus aureus during manganese starvation. Nucleic Acids Res, 47, 9871–9887.

23. Westermann, A.J., Venturini, E., Sellin, M.E., Forstner, K.U., Hardt, W.D., and Vogel, J. (2019) The Major RNA-Binding Protein ProQ Impacts Virulence Gene Expression in Salmonella enterica Serovar Typhimurium. mBio, 10.

24. Kim, K., Palmer, A.D., Vanderpool, C.K., and Slauch, J.M. (2019) The Small RNA PinT Contributes to PhoP-Mediated Regulation of the Salmonella Pathogenicity Island 1 Type III Secretion System in Salmonella enterica Serovar Typhimurium. J Bacteriol, 201.

25. Arnvig, K.B., and Young, D.B. (2009) Identification of small RNAs in Mycobacterium tuberculosis. Molecular Microbiology, 73, 397–408.

26. Arnvig, K.B., Comas, I., Thomson, N.R., Houghton, J., Boshoff, H.I., Croucher, N.J., Rose, G., Perkins, T.T., Parkhill, J., Dougan, G., et al. (2011) Sequence-based analysis uncovers an abundance of non-coding RNA in the total transcriptome of Mycobacterium tuberculosis. PLoS Pathog, 7, e1002342.

27. Miotto, P., Forti, F., Ambrosi, A., Pellin, D., Veiga, D.F., Balazsi, G., Gennaro, M.L., Di Serio, C., Ghisotti, D., and Cirillo, D.M. (2012) Genome-wide discovery of small RNAs in Mycobacterium tuberculosis. PLoS One, 7, e51950.

28. Li, S.-K., Ng, P.K.-S., Qin, H., Lau, J.K.-Y., Lau, J.P.-Y., Tsui, S.K.-W., Chan, T.-F. and Lau, T.C.-K. (2012) Identification of small RNAs in Mycobacterium smegmatis using heterologous Hfq. RNA.

29. Ami, V.K.G., Balasubramanian, R., and Hegde, S.R. (2020) Genome-wide identification of the context-dependent sRNA expression in Mycobacterium tuberculosis. BMC Genomics, 21, 167.

30. Hnilicová, J., Jirát Matějčková, J., Siková, M., Pospíšil, J., Halada, P., Pánek, J., and Krásny, L. (2014) Ms1, a novel sRNA interacting with the RNA polymerase core in mycobacteria. Nucleic Acids Research.

31. Girardin, R.C., and McDonough, K.A. (2020) Small RNA Mcr11 requires the transcription factor AbmR for stable expression and regulates genes involved in the central metabolism of Mycobacterium tuberculosis. Mol Microbiol, 113, 504–520.

32. Solans, L., Gonzalo-Asensio, J., Sala, C., Benjak, A., Uplekar, S., Rougemont, J., Guilhot, C., Malaga, W., Martín, C., and Cole, S.T. (2014) The PhoP-Dependent ncRNA Mcr7 Modulates the TAT Secretion System in Mycobacterium tuberculosis. PLoS Pathogens, 10, e1004183.

33. Gerrick, E.R., Barbier, T., Chase, M.R., Xu, R., Francois, J., Lin, V.H., Szucs, M.J., Rock, J.M., Ahmad, R., Tjaden, B., et al. (2018) Small RNA profiling in Mycobacterium tuberculosis identifies MrsI as necessary for an anticipatory iron sparing response. Proc Natl Acad Sci U S A, 115, 6464–6469.

34. Salina, E.G., Grigorov, A., Skvortsova, Y., Majorov, K., Bychenko, O., Ostrik, A., Logunova, N., Ignatov, D., Kaprelyants, A., Apt, A., et al. (2019) MTS1338, A Small Mycobacterium tuberculosis RNA, Regulates Transcriptional Shifts Consistent With Bacterial Adaptation for Entering Into Dormancy and Survival Within Host Macrophages. Front Cell Infect Microbiol, 9, 405.

35. Singh, S., Nirban, R., and Dutta, T. (2021) MTS1338 in Mycobacterium tuberculosis promotes detoxification of reactive oxygen species under oxidative stress. Tuberculosis (Edinb), 131, 102142.

36. Mai, J., Rao, C., Watt, J., Sun, X., Lin, C., Zhang, L., and Liu, J. (2019) Mycobacterium tuberculosis 6C sRNA binds multiple mRNA targets via C-rich loops independent of RNA chaperones. Nucleic Acids Res, 47, 4292–4307.

37. Miranda-CasoLuengo, A.A., Staunton, P.M., Dinan, A.M., Lohan, A.J., and Loftus, B.J. (2016) Functional characterization of the Mycobacterium abscessus genome coupled with condition specific transcriptomics reveals conserved molecular strategies for host adaptation and persistence. BMC Genomics, 17, 553.

38. Foreman, M., Gershoni, M., and Barkan, D. (2020) A Simplified and Efficient Method for Himar-1 Transposon Sequencing in Bacteria, Demonstrated by Creation and Analysis of a Saturated Transposon-Mutant Library in Mycobacterium abscessus. mSystems, 5.

39. Martini, M.C., Zhou, Y., Sun, H., and Shell, S.S. (2019) Defining the Transcriptional and Post-transcriptional Landscapes of Mycobacterium smegmatis in Aerobic Growth and Hypoxia. Front Microbiol, 10, 591.

40. Cortes, T., Schubert, O.T., Rose, G., Arnvig, K.B., Comas, I., Aebersold, R., and Young, D.B. (2013) Genome-wide Mapping of Transcriptional Start Sites Defines an Extensive Leaderless Transcriptome in Mycobacterium tuberculosis. Cell reports.

41. Shell, S.S., Wang, J., Lapierre, P., Mir, M., Chase, M.R., Pyle, M.M., Gawande, R., Ahmad, R., Sarracino, D.A., Ioerger, T.R., et al. (2015) Leaderless Transcripts and Small Proteins Are Common Features of the Mycobacterial Translational Landscape. PLoS Genetics, 11, e1005641.

42. Meir, M., Grosfeld, T., and Barkan, D. (2018) Establishment and Validation of Galleria mellonella as a Novel Model Organism To Study Mycobacterium abscessus Infection, Pathogenesis, and Treatment. Antimicrob Agents Chemother, 62.

43. Rao, V., Gao, F., Chen, B., Jacobs, W.R., Jr. and Glickman, M.S. (2006) Trans-cyclopropanation of mycolic acids on trehalose dimycolate suppresses Mycobacterium tuberculosis -induced inflammation and virulence. J Clin Invest, 116, 1660–1667.

44. Riva, C., Tortoli, E., Cugnata, F., Sanvito, F., Esposito, A., Rossi, M., Colarieti, A., Canu, T., Cigana, C., Bragonzi, A., et al. (2020) A New Model of Chronic Mycobacterium abscessus Lung Infection in Immunocompetent Mice. Int J Mol Sci, 21.

45. Lore, N.I., Saliu, F., Spitaleri, A., Schafle, D., Nicola, F., Cirillo, D.M., and Sander, P. (2022) The aminoglycoside modifying enzyme Eis2 represents a new potential in vivo target for reducing antimicrobial drug resistance in Mycobacterium abscessus complex. Eur Respir J.

46. Sharp, J.S., and Bechhofer, D.H. (2003) Effect of translational signals on mRNA decay in Bacillus subtilis. Journal of Bacteriology, 185, 5372–5379.

47. Proshkin, S., Rahmouni, A.R., Mironov, A., and Nudler, E. (2010) Cooperation between translating ribosomes and RNA polymerase in transcription elongation. Science, 328, 504–508.

48. Hambraeus, G., Karhumaa, K., and Rutberg, B. (2002) A 5’ stem-loop and ribosome binding but not translation are important for the stability of Bacillus subtilis aprE leader mRNA. Microbiology (Reading, England), 148, 1795–1803.

49. Jurgen, B., Schweder, T., and Hecker, M. (1998) The stability of mRNA from the gsiB gene of Bacillus subtilis is dependent on the presence of a strong ribosome binding site. Mol Gen Genet, 258, 538–545.

50. Richardson, J.P. (1991) Preventing the synthesis of unused transcripts by Rho factor. Cell, 64, 1047–1049.

51. Pato, M.L., Bennett, P.M., and von Meyenburg, K. (1973) Messenger ribonucleic acid synthesis and degradation in Escherichia coli during inhibition of translation. Journal of Bacteriology, 116, 710–718.

52. Varmus, H.E., Perlman, R.L., and Pastan, I. (1971) Regulation of lac transcription in antibiotic-treated E. coli. Nat New Biol, 230, 41–44.

53. Deana, A., and Belasco, J.G. (2005) Lost in translation: the influence of ribosomes on bacterial mRNA decay. Genes & Development, 19, 2526–2533.

54. Redondo, N., Mok, S., Montgomery, L., Flanagan, P.R., McNamara, E., Smyth, E.G., O’Sullivan, N., Schaffer, K., Rogers, T.R., and Fitzgibbon, M.M. (2020) Genomic Analysis of Mycobacterium abscessus Complex Isolates Collected in Ireland between 2006 and 2017. J Clin Microbiol, 58.

55. Ripoll, F., Deshayes, C., Pasek, S., Laval, F., Beretti, J.L., Biet, F., Risler, J.L., Daffe, M., Etienne, G., Gaillard, J.L., et al. (2007) Genomics of glycopeptidolipid biosynthesis in Mycobacterium abscessus and M. chelonae. BMC Genomics, 8, 114.

56. Budell, W.C., Germain, G.A., Janisch, N., McKie-Krisberg, Z., Jayaprakash, A.D., Resnick, A.E., and Quadri, L.E.N. (2020) Transposon mutagenesis in Mycobacterium kansasii links a small RNA gene to colony morphology and biofilm formation and identifies 9,885 intragenic insertions that do not compromise colony outgrowth. Microbiologyopen, 9, e988.

57. Clary, G., Sasindran, S.J., Nesbitt, N., Mason, L., Cole, S., Azad, A., McCoy, K., Schlesinger, L.S., and Hall-Stoodley, L. (2018) Mycobacterium abscessus Smooth and Rough Morphotypes Form Antimicrobial-Tolerant Biofilm Phenotypes but Are Killed by Acetic Acid. Antimicrob Agents Chemother, 62.

58. Barkan, D., Rao, V., Sukenick, G.D., and Glickman, M.S. (2010) Redundant function of cmaA2 and mmaA2 in Mycobacterium tuberculosis cis cyclopropanation of oxygenated mycolates. J Bacteriol, 192, 3661–3668.

59. Vargas-Blanco, D.A., Zhou, Y., Zamalloa, L.G., Antonelli, T., and Shell, S.S. (2019) mRNA Degradation Rates Are Coupled to Metabolic Status in Mycobacterium smegmatis. MBio, 10.

60. Culviner, P.H., Guegler, C.K., and Laub, M.T. (2020) A Simple, Cost-Effective, and Robust Method for rRNA Depletion in RNA-Sequencing Studies. mBio, 11.

61. Culviner, P.H., and Laub, M.T. (2018) Global Analysis of the E. coli Toxin MazF Reveals Widespread Cleavage of mRNA and the Inhibition of rRNA Maturation and Ribosome Biogenesis. Mol Cell, 70, 868–880 e810.

62. Martin, M. (2011) Cutadapt Removes Adapter Sequences From High-Throughput Sequencing Reads. EMBnet.journal, 17, 10–12.

63. Li, H., and Durbin, R. (2009) Fast and accurate short read alignment with Burrows-Wheeler transform. Bioinformatics, 25, 1754–1760.

64. Liao, Y., Smyth, G.K., and Shi, W. (2014) featureCounts: an efficient general purpose program for assigning sequence reads to genomic features. Bioinformatics, 30, 923–930.

65. Love, M.I., Huber, W., and Anders, S. (2014) Moderated estimation of fold change and dispersion for RNA-seq data with DESeq2. Genome Biol, 15, 550.

66. van Kessel, J.C., and Hatfull, G.F. (2007) Recombineering in Mycobacterium tuberculosis. Nat Methods, 4, 147–152.

67. Tufariello, J.M., Malek, A.A., Vilchèze, C., Cole, L.E., Ratner, H.K., González, P.A., Jain, P., Hatfull, G.F., Larsen, M.H., and Jacobs, W.R. (2014) Enhanced Specialized Transduction Using Recombineering in Mycobacterium tuberculosis. mBio, 5.

68. Facchini, M., De Fino, I., Riva, C., and Bragonzi, A. (2014) Long term chronic Pseudomonas aeruginosa airway infection in mice. J Vis Exp.

69. Meir, M., Bifani, P., and Barkan, D. (2018) The addition of avibactam renders piperacillin an effective treatment for Mycobacterium abscessus infection in an in vivo model. Antimicrob Resist Infect Control, 7, 151.

70. Li, H., Handsaker, B., Wysoker, A., Fennell, T., Ruan, J., Homer, N., Marth, G., Abecasis, G., Durbin, R., and Genome Project Data Processing, S. (2009) The Sequence Alignment/Map format and SAMtools. Bioinformatics, 25, 2078–2079.

71. Carver, T., Harris, S.R., Berriman, M., Parkhill, J., and McQuillan, J.A. (2012) Artemis: an integrated platform for visualization and analysis of high-throughput sequence-based experimental data. Bioinformatics (Oxford, England), 28, 464–469.

